# Stepwise design of pseudosymmetric protein hetero-oligomers

**DOI:** 10.1101/2023.04.07.535760

**Authors:** Ryan D. Kibler, Sangmin Lee, Madison A. Kennedy, Basile I. M. Wicky, Stella M. Lai, Marius M. Kostelic, Xinting Li, Cameron M. Chow, Lauren Carter, Vicki H. Wysocki, Barry L. Stoddard, David Baker

**Affiliations:** Department of Biochemistry, University of Washington, Seattle, WA 98195, USA; Institute for Protein Design, University of Washington, Seattle, WA 98195, USA; Howard Hughes Medical Institute, University of Washington, Seattle, WA 98195, USA; Division of Basic Sciences, Fred Hutchinson Cancer Center, Seattle, WA 98006, USA; Department of Chemistry and Biochemistry, The Ohio State University, Columbus, OH 43210, USA; Resource for Native Mass Spectrometry Guided Structural Biology, The Ohio State University, Columbus, OH 43210, USA

## Abstract

Pseudosymmetric hetero-oligomers with three or more unique subunits with overall structural (but not sequence) symmetry play key roles in biology, and systematic approaches for generating such proteins *de novo* would provide new routes to controlling cell signaling and designing complex protein materials. However, the *de novo* design of protein hetero-oligomers with three or more distinct chains with nearly identical structures is a challenging problem because it requires the accurate design of multiple protein-protein interfaces simultaneously. Here, we describe a divide-and-conquer approach that breaks the multiple-interface design challenge into a set of more tractable symmetric single-interface redesign problems, followed by structural recombination of the validated homo-oligomers into pseudosymmetric hetero-oligomers. Starting from *de novo* designed circular homo-oligomers composed of 9 or 24 tandemly repeated units, we redesigned the inter-subunit interfaces to generate 15 new homo-oligomers and recombined them to make 17 new hetero-oligomers, including ABC heterotrimers, A2B2 heterotetramers, and A3B3 and A2B2C2 heterohexamers which assemble with high structural specificity. The symmetric homo-oligomers and pseudosymmetric hetero-oligomers generated for each system share a common backbone, and hence are ideal building blocks for generating and functionalizing larger symmetric assemblies.

**Significance Statement:** Protein oligomers composed of multiple unique subunits are versatile building blocks for creating functional materials and controlling biological processes. However, designing robust hetero-oligomers with distinct subunits and precise structural symmetry remains a major challenge. Here, we present a general strategy for designing such complexes by breaking down the problem into simpler steps by first symmetrically re-designing the interfaces of homo-oligomeric proteins, and then recombining validated variants to form pseudosymmetric hetero-oligomers. Using this method, we generated 17 hetero-oligomers with up to three unique subunits that assemble with high specificity. Our approach can be extended to create a wide range of pseudosymmetric assemblies for manipulating cellular signaling and as building blocks for advanced protein materials. These pseudosymmeteric heterotrimers have already enabled the construction of a set of massive nanocages, including a T=4 icosahedral nanocage with a 70 nm diameter and 240 subunits.^1^

## Main text

Pseudosymmetric hetero-oligomers in Nature likely evolved from homo-oligomers composed of identical subunits which, following gene duplication, underwent divergence and specialization leading to structures where each subunit is structurally similar but can have different specialized functions^2,3^ (Fig. 1a). Specialization of binding surfaces can create hetero-oligomeric assembly hubs where each subunit can recruit a unique set of auxiliary proteins. For example, the eukaryotic pseudosymmetric heteroheptameric and -hexameric complexes of Sm and LSm proteins, which share a common ancestor with the bacterial homohexamer Hfq^4,5^, mediate a wide network of RNA-protein interactions via specific combinations of protomers^4,6^. Another example, PCNA, acts as a homotrimeric DNA-modifying protein recruitment hub in eukaryotes, but forms pseudosymmetric heterotrimers in archaea with unique protein binding sites on each subunit and a different assembly pathway^7^. More generally, evolution has used pseudosymmetrization as a route to enhance the functionality of originally homo-oligomeric assemblies. As in Nature, pseudosymmetrization of oligomers could provide a route to enhancing the functionality and complexity of designed protein assemblies. In particular, as described in the accompanying manuscript (Lee et al), pseudosymmetrization provides a route from designed icosahedral assemblies with triangulation number = 1 and 60 identical subunits to more complex higher triangulation number assemblies with multiple distinct subunits and much larger interior volumes for packaging cargoes. However, while there has been considerable progress with *de novo* design of protein homo- and hetero-oligomers^8–18^, with the exception of heterodimers and coiled coil heterotrimers, there are no systematic routes to pseudosymmetric structures with nearly identical protomer backbones but different sequences. This is a challenging problem because, for a pseudosymmetric cyclic n-mer with n distinct chains in the target ground state oligomer, there are on the order of n^n-1^ off target assemblies, all with similar or identical backbone geometry. Therefore, the specificity for the intended heteromeric complex must come from the amino acid sidechain identities.

**Fig. 1.**
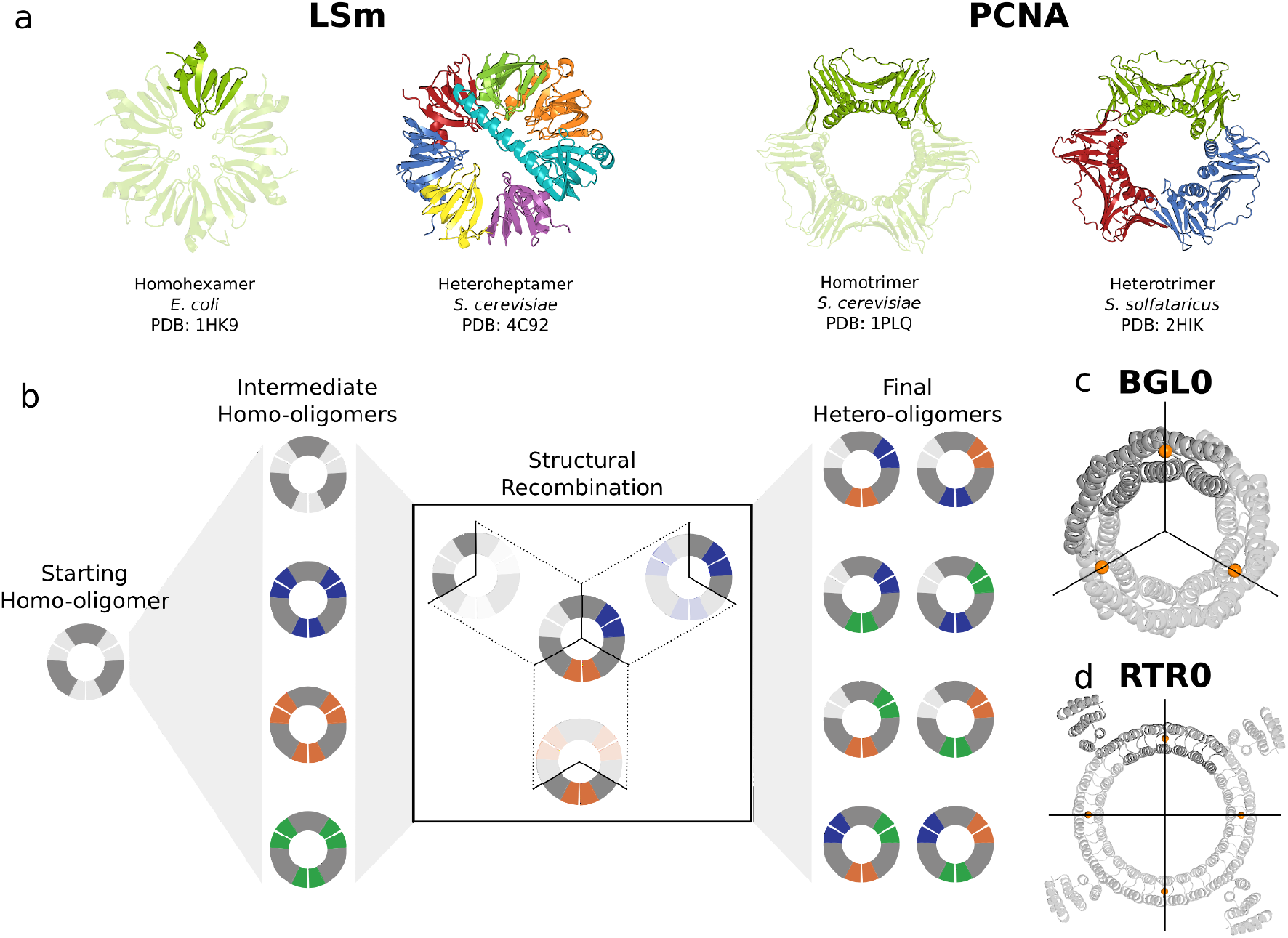
Overview of pseudosymmetic recombination design approach. **(a)** Two examples of natural pseudosymmetric hetero-oligomers (each pair, right) which likely arose from symmetric ancestors, resembling the paired symmetric paralogs (each pair, left), through gene duplication and diversification. Left, LSm complexes from *Escherichia coli* (Hfq) and *Saccharomyces cerevisiae*. Right, PCNA from *S. cerevisiae* and *Sulfolbus solfataricus*. **(b)** Diagram of our stepwise hetero-oligomer creation strategy; variant homo-oligomeric interfaces are generated from a common structure first, followed by junction splicing to assemble theminto hetero-oligomers. Cartoons represent trimers with colored interfaces that match up (separated by a small white gap in the cartoon) and have unique sequences according to their color. Interfaces on the same chain are connected via a homology region in dark gray which has the same sequence and structure across all protomer variants. In this strategy, the interface of an existing homo-oligomer (left; a homotrimer in this example) is redesigned symmetrically and the junction regions remain unchanged. Homo-oligomers that assemble correctly (left column) are cut at the same point within their common homology regions and then spliced back together, making two-chain hybrids with two different interfaces (Fig. S1b) which assemble into heterotrimers (right column). It is possible to create two different heterotrimers with the same interfaces by rearranging the order of the interfaces (Fig. S1c). **(c,d)** The two test scaffolds used in this work, the homotrimer BGL0 (c) and the homotetramer RTR0 (d). Each is bisected by black crosshairs which pass through the orange sphere on each chain which indicates the location of the residue (Arg82 and Leu98 for BGL0 and RTR0, respectively) within the junction region where splicing will occur (Fig. S1b).

We set out to develop a general *de novo* design approach for creating pseudosymmetric hetero-oligomers (Fig. 1b) inspired by the likely evolutionary route from symmetric homo-oligomers to hetero-oligomers (Fig. 1a). We focused on architectures in which the interfaces between different subunits are structurally disjoint; in the cyclic case these correspond to circular ring-like assemblies with each subunit forming part of the arc of the ring (see Fig. 1c,d). Such architectures have the advantage that each interface can be validated independently in the context of a homo-oligomer that can be experimentally characterized on its own. We aimed to make each of these inter-subunit interfaces distinct by incorporating different hydrogen bonding networks (“HBNets”); because there is a large energetic penalty disfavoring burial of polar groups, interfaces with extensive hydrogen bond networks have higher specificity than purely hydrophobic interfaces, with reduced off-target interactions^10^. HBNets have been used previously to create a large number of orthogonal heterodimeric interfaces on similarly-shaped helical backbones^11^.

We developed a stepwise design strategy for generating hetero-oligomers with distinct hydrogen bond networks at each subunit-subunit interface (Fig. 1b). First, we redesign a homo-oligomeric scaffold and experimentally characterize multiple variants, each with distinct HBNets at the inter-subunit interface^10,11^. We increase the variety of HBNets that can be incorporated by introducing small backbone perturbations in the secondary structure elements present at the interface^19^. In the second step, we recombine experimentally validated homo-oligomers in a manner analogous to DNA recombination to generate pseudosymmetric hetero-oligomers. This “divide and conquer” approach avoids the highly error-prone approach of designing multiple interfaces simultaneously (“one-shot design”) and also generates a large number of hetero-oligomers through the recombination of a small set of homo-oligomers (Fig. S1a).

### Pseudosymmetrizing a computationally designed 9 repeat toroid

We chose as a first model system a circular Tandem Repeat Protein or “cTRP” composed of three identical subunits forming a ring (Fig. 1c, tcTRP9_sub3 from ref ^20^ referred to here as BGL0). Because Rosetta HBNet placement is very sensitive to small backbone displacements^19^, we enhanced HBNet sampling without drastically altering the interface by generating small backbone variations at the interface using normal modes (“NM”) relaxation^21^ restricted to the interface (NM design set) or by complete backbone resampling of the C-terminal helix which is on the outer ring of helices and forms the HBNet interface in conjunction with two inner ring helices. In the latter resampling, we varied the internal geometry of the helix using the Crick alpha-helix parameters^22^ starting from the closest fit parameters to the C-terminal helix and varying r_0_, ω_0_, ω_1_, φ_0_, φ_1_, and Δz, and then rejoined the helix to the C-terminus (HR-C design set) or, alternatively, attached it anew to the N-terminus (HR-N design set). The HR-N design set was expected to be orthogonal to the HR-C and NM designs due to steric hindrance imposed by mismatching terminal helices (clashing in one case, a large gap with reduced hydrophobic contacts in the other).

We obtained synthetic genes encoding 20 redesigns of BGL0 (named BGL01-20, Table S1) and produced the proteins in *E. coli*. Out of the 20 tested, 10 (NM: 3/4; HR-N: 3/8; HR-C: 4/8) were trimeric according to size exclusion chromatography (SEC) (Fig. 2b, Fig. S2), native mass spectrometry (nMS)^23,24^ (Fig. 2c, Fig. S3), and SEC-Multi-Angle Light Scattering (SEC-MALS) (Fig. S4); the most common failure mode was polydisperse or aggregated SEC traces (Fig. S2). Small-angle X-ray scattering (SAXS)^25,26^ profiles (Fig. 2d, Fig. S5) and negative-stain electron microscopy (nsEM) 2D class averages (Fig. 2e) of these designs are consistent with the design models.

**Figure 2:**
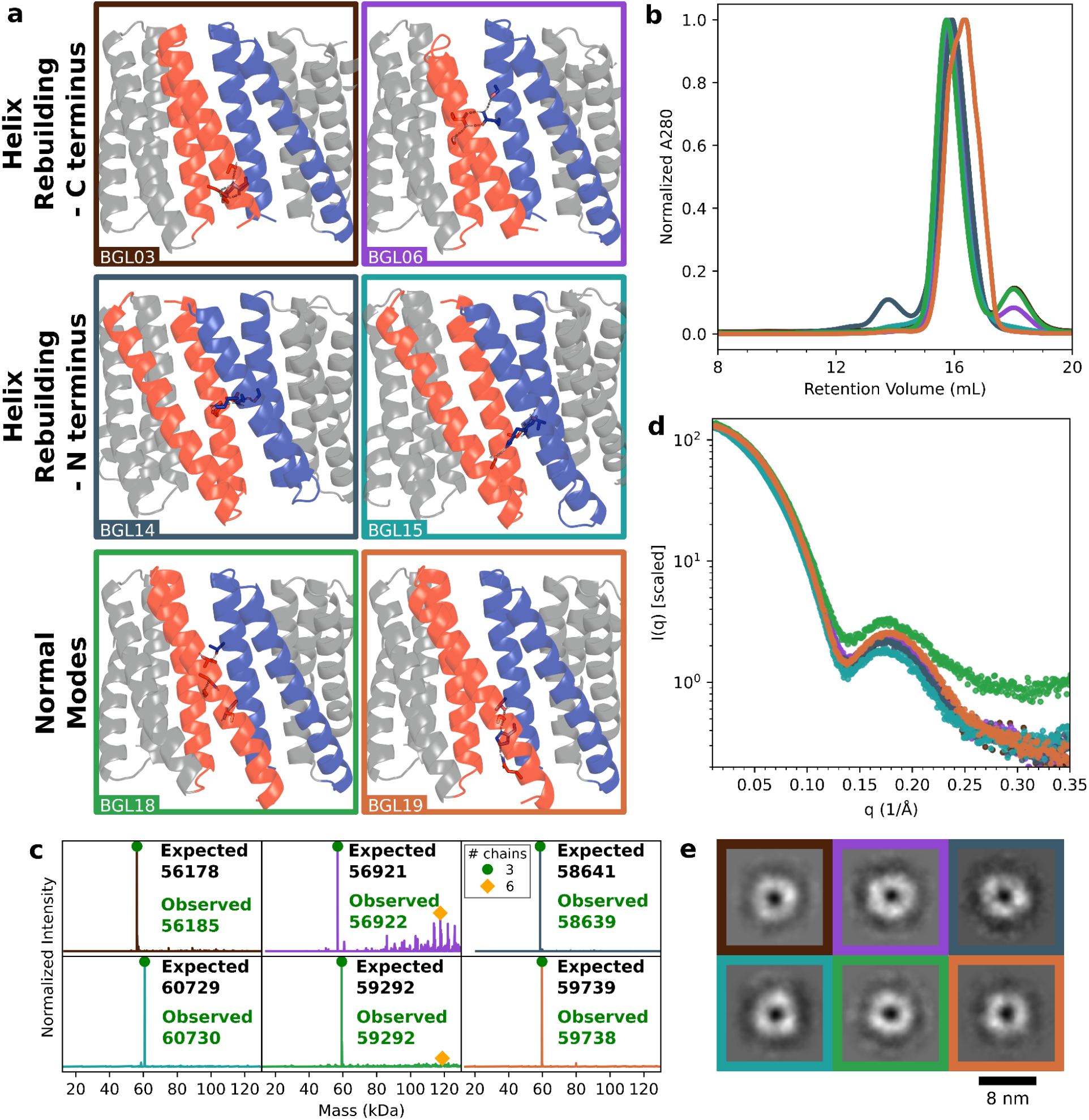
Characterization of BGL homotrimers. **(a)** Cartoon representation of 6 representative successful redesigns of BGL0, with two from each design set: Helix Rebuilding + C terminal attachment (HR-C), Helix Rebuilding + N terminal attachment (HR-N) and Normal Modes (NM) relaxation. Blue and red colored backbones highlight N- and C-terminal two-helix segments comprising the interface which was redesigned, and newly designed hydrogen bond networks (HBNets) are shown in sticks with the hydrogen bonds in dashed lines. Gray backbones are supporting structures or other copies of the interface. Design names in the bottom left corner of each box and box outline colors are used consistently throughout. Characterization of the six examples from (**a**) by (b) SEC, (c) nMS, (d) SAXS, and (e) nsEM. **(b)** Overlay of A280 normalized SEC traces. Samples were previously purified by SEC to remove any soluble aggregate. SEC traces of all 20 designs are provided in Supplemental Figure 2. **(c)** Individual mass deconvoluted nMS data of TEV-cleaved samples with all peaks that match the expected mass for each homotrimer are annotated with green circles, and homohexamers (likely to be dimers of trimers) are annotated with orange diamonds. Trace color corresponds to sample ID. See Table S7 for complete data, including m/z spectra. **(d)** Overlay of SAXS data. Data were scaled appropriately to align at q=0.01 for the purpose of displaying similarity of profile. Radii of gyration calculated from the SAXS profiles match the design models within 5.7Å. See Supplemental Figure 5. **(e)** Individual 2D class averages of nsEM data collected on each sample. Outline color corresponds to sample ID.

We obtained crystal structures of BGL06 from the HR-C set (Fig. 3a), BGL14 and BGL15 from the HR-N set (Fig. 3b,c), and BGL18 from the NM set (Fig. 3d). At the backbone level, the crystal structures show good agreement with the design models (1.2 Å, 1.3 Å, 0.7 Å, and 1.4 Å CA-RMSD, respectively), as well as to the parental BGL0 crystal structure (TM-scores 0.94, 0.82, 0.81, and 0.97, respectively, to PDB 6XR2). Sidechains at the interfaces of BGL06, BGL14, and BGL18 are well enough resolved to determine that HBNets in all three structures are formed correctly. BGL06 has an interchain bidentate HBNet involving Gln14 and Gln146, each backed up with second shell interactions from Ser18 and Ser125, respectively (Fig. 3a). The string of hydrogen bonds in the BGL14 interface alternates between chains, with a fully satisfied Asn13 in the very middle which donates a hydrogen bond to His153. BGL15 has three copies of the ring in the unit cell, but the low resolution of the dataset (4.0 Å) obscured details about the HBNet residue sidechains. BGL18 has two copies of the ring in the unit cell and all six interfaces were resolved and highly similar to each other. The other copy of the trimer in the unit cell is 1.6Å CA-RMSD to the design model. It has a generally polar interface with a meandering HBNet spanning Asn10, Thr17, Asn126 (which may be donating a hydrogen bond to the backbone oxygen of Asn10), Thr152, Ser155, and Ser159.

**Figure 3.**
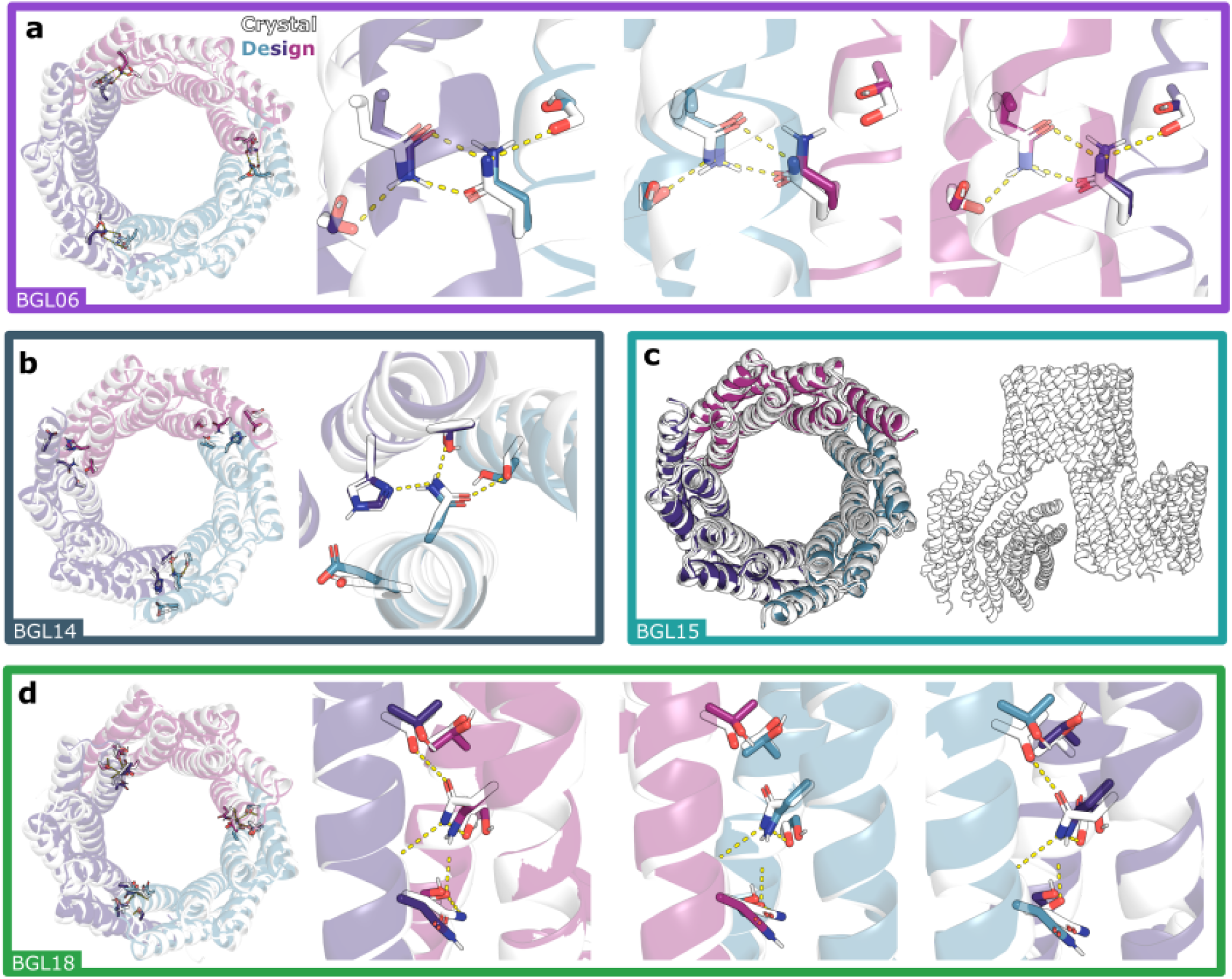
Crystal structures of BGL homotrimers. **(a,b,d)** Crystal structures (white) of four different BGL homotrimers are superimposed onto the design models which are colored in teal, purple, and magenta. (left) The top-down view of the full ring with HBNets highlighted and visible through transparent cartoon models. The backbones of all crystal structures match well with the design models (<1.4 Å RMSD over all Cα atoms). (Right) Close up view(s) of each HBNet well enough resolved to build full side chains. **(a)** BGL06 (2.1 Å resolution; RMSD = 1.2 Å over all Cα atoms; PDB: 8E0L) and **(d)** BGL18 (3.0 Å resolution; RMSD = 1.4 Å over all Cα atoms; PDB: 8E0N) were both well enough resolved to model all three interfaces. BGL18 contained two copies of the full trimer in the crystal unit cell, which are structurally similar to each other (RMSD = 0.3 Å over all Cα atoms between copies). **(b)** One interface of BGL14 (3.0 Å resolution; RMSD = 1.3 Å over all Cα atoms; PDB: 8E12) was well resolved. **(c)** BGL15 (4.0Å resolution, PDB: 8E0M) had three copies of the trimer in the unit cell and all three were close to the design model (0.63 Å, 0.64 Å, and 0.66 Å RMSD over Cα atoms). See Table S2 for crystallographic details.

We constructed multiple pseudosymmetric trimers by selecting 3 homotrimers at a time from the 11 validated designs (10 redesigns + the original) and recombining them as illustrated in Fig. 1b and Fig. S1. Each subunit was split in the middle at Arg82, which is far from either interface. Complementary pairs were then reconnected at the splicing junction to form hybrid protomers (Fig. S1b) which in principle can assemble into a heterotrimer. Two distinct heterotrimers can be generated from each selected set of homotrimeric interfaces (*a,b*, and *c)* by changing the order of recombination along the circle: (*a*-*b*; *b*-*c*; *c*-*a*) and (*a*-*c*; *c*-*b*; *b*-*a*) (Fig. S1c). This procedure can generate considerable diversity; the number of possible hetero-n-mers grows very quickly with the number of validated homo-n-mers from which the interfaces are extracted (Fig. S1a): there are

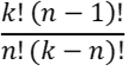

ways to combine *k* interfaces into a hetero-n-mer with n different interfaces. With k=11 interfaces to choose from for a heterotrimer (n=3), there are 330 possible heterotrimeric variants that could theoretically be generated.

We tested two different strategies for choosing interfaces for recombination. First, to attempt to maximize binding specificity, we constructed a set of 6 recombinants (called “Set 1”) with one interface from each of HR-N, HR-C, and NM. Second, to generate heterotrimers with nearly identical subunit backbone structures, we recombined the original design BGL0 with validated homo-oligomers from the less-perturbed NM set (BGL17, BGL18, and BGL19). From the four homo-oligomers we enumerated all 8 possible combinations (called “Set 2”).

Both sets of designed heterotrimers (total 14,see Table S3) were ordered as tricistronic constructs with TEVp-cleavable GFP or miniprotein^27^ tags on two of the chains to differentiate their masses by SDS-polyacrylamide gel electrophoresis (SDS-PAGE) and nMS, and the third chain having a His6-tag for affinity co-purification (Fig. 4a). We found that 11/14 co-purified with three distinct bands on SDS-PAGE, with 9 showing roughly stoichiometric ratios by gel densitometry (Fig. 4b, Fig. S6). The SEC profiles of 8 (2 from Set 1 and 6 from Set 2) were largely monodisperse and had the expected retention volume (Fig. 4c, Fig. S7). nMS analyses of these 8 designs showed that each was the intended ABC species (consisting of the three unique chains: A, B, and C) without any of the many possible off-target combinations of subunits (AAB, BBB, etc; Fig. 4d,f; Fig. S8). nsEM and SAXS data confirmed that these eight heterotrimers retained the original toroidal shape (Fig. S9, Fig. S10). Each subunit of a heterotrimer from Set 1 has a different number of helices due to the use of interfaces from the HR-N set with interfaces from the HR-C or NM sets. To make them more like the subunits of heterotrimers from Set 2, which have the same number of helices in each subunit, we deleted the loop of the N-terminal outer helix in the HR-N-derived interface and reconnected it at the C-terminus of the next protomer (essentially converting the HR-N interface to HR-C); 2/3 such rearrangements remained clean heterotrimers (Fig. S11), which brought the number of validated hetBGLs to 10.

**Figure 4:**
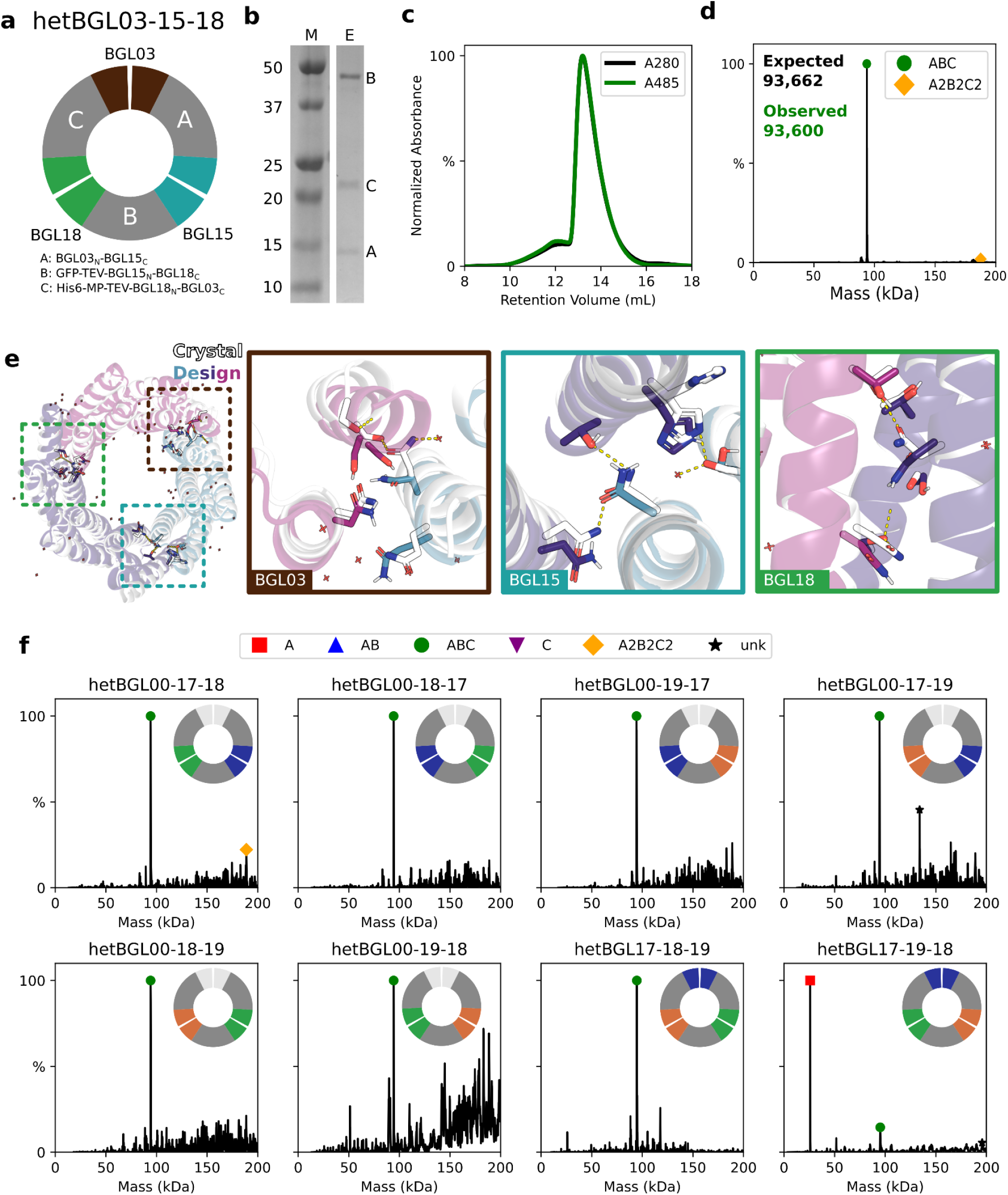
Characterization of hetBGL heterotrimers. **(a-e)** Results of hetBGL03-15-18. **(a)** (top) Schematic showing the result of recombination between BGL03, BGL15, and BGL18. (bottom) The expression constructs for each chain as expressed tricistronically. “GFP” is superfolderGFP, included to add a large amount of mass and provide an additional spectroscopic identification, and “MP” is EHEE_rd2_0005^27^, a stable and small protein included to add a small amount of mass. **(b)** SDS-PAGE of IMAC elution. M: protein ladder. E: IMAC elution. Protein size affects band staining, so it is difficult to judge stoichiometry from the band darkness/size, but the correct trend in stain density is observed for equal stoichiometry. **(c)** SEC trace overlaying normalized absorbances at A280 and A485. **(d)** Mass deconvoluted nMS data showing the intended major species (green circle, ABC) and a minor species with twice the mass, likely to be a dimer of trimers (orange diamond, A2B2C2). See Table S7 for complete data, including m/z spectra. **(e)** (left) Crystal structure in white (2.1 Å resolution, PDB: 8E0O) and design model in teal, purple, and magenta overlaid to show global agreement (CA-RMSD: 1.2 Å, TM-score: 0.96). Dashed boxes indicate approximate locations of zoomed-in views of the interfaces derived from BGL03, BGL15, and BGL18, showing side chains of HBNet residues and their inferred hydrogen bonds as yellow dashes. **(f)** Individual mass deconvoluted nMS data for all hetBGL Set 2 samples. See Table S7 for complete data, including m/z spectra.

We solved the crystal structure of hetBGL03-15-18, a heterotrimer from Set 1 composed of the interfaces extracted from BGL03, BGL15, and BGL18, at 2.1 Å resolution (Fig. 4e). The structure superimposes to the design model with 1.2 Å CA-RMSD. The three distinct subunits have an average pairwise sequence identity of 62.3% but high structural similarity (aligned CA-RMSD of 1.5 Å), showing that this method can produce highly (pseudo)symmetric hetero-oligomers. The BGL18-derived HBNet is very similar to that in the homotrimer crystal structure, and the HBNet of BGL15, which was not resolved in the homotrimer crystal structure, is confirmed to be in the correct conformation, with the addition of a hydrogen bond between Ser140 and a crystallographic water. The previously unsolved BGL03 interface differs from the design model due to fraying of the C-terminal outer helix likely due to reduction of hydrophobic packing stemming from polar HBNet residues placed too close to the C-terminus; placement of HBNet residues too close to the termini was avoided in future designs.

### Pseudosymmetrizing a computationally designed 24 repeat toroid

Next we tested whether our hierarchical pseudo-symmetrization method could be generalized to another homo-oligomeric system. We selected a designed tetrameric toroid with 24 internal repeats separated into four protomers of 6 repeats each (Fig. 1d, C4_1 from ref^28^ referred to here as RTR0). Since there are multiple ways of factoring 24 repeats into protomers, this system can be readily redesigned to generate C3, C4, C6, and C12 assemblies (Fig. S12a), offering multiple possibilities for hetero-oligomerization (trimers: ABC; tetramers: A2B2, ABCD; hexamers: A3B3, A2B2C2, ABCDEF; etc). Based on the success of the NM set of the BGL family, we only used NM perturbation with the RTR family. HBNets and the surrounding sequences at the interfaces were designed similarly to BGL0, with the exception of avoiding the addition of HBNet residues near the ends of the N-terminal helix. We also installed disulfide bonds at diverse positions,^29^ since interface disulfides were required for the original toroid to form tetramers and higher-order symmetries^30^; the disulfides could also provide an additional level of specificity (Fig. 5a).

**Figure 5:**
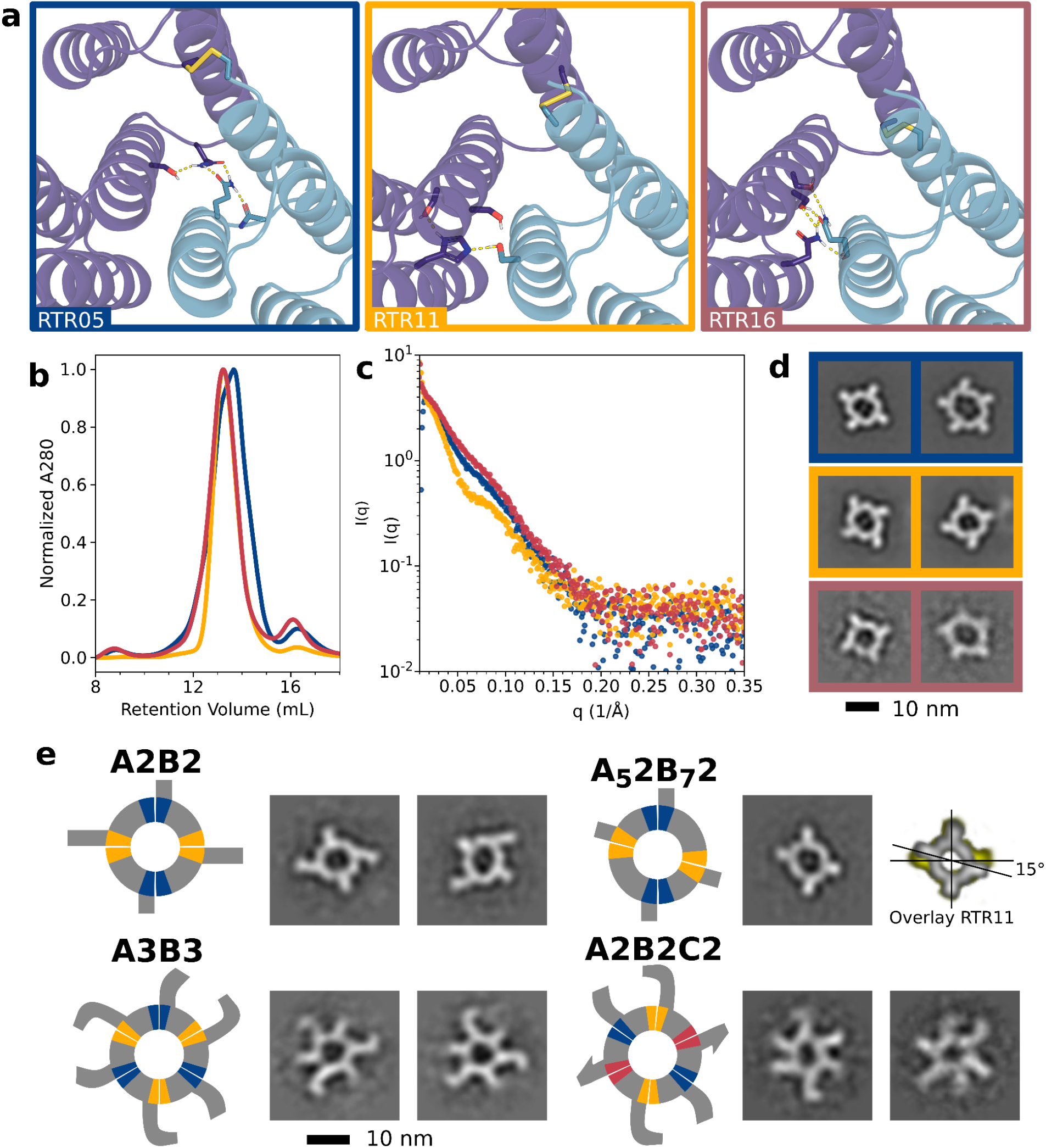
Characterization of RTR homotetramers and hetero-oligomers. **(a)** Cartoon representation of 3 representative successful redesigns of RTR0. Blue and purple colored backbones represent N- and C-terminal interfaces, respectively. HBNets and disulfide bonds are shown in sticks with hydrogen bonds in dashed lines. Design names (bottom left corner of each box) and box outline colors are used consistently throughout to represent the same interfaces. Characterization of the 3 examples from (a) by (b) SEC, (c) nMS, (d) SAXS, and (e) nsEM. **(b)** Overlay of A280 normalized SEC traces. SEC traces of all 24 designs are provided in Supplemental Figure 12. **(c)** Overlay of SAXS data. Radii of gyration calculated from the SAXS profiles match the design models within 7.2 Å. **(d)** Individual 2D class averages of nsEM data collected on each sample. Outline color corresponds to sample ID. **(e)** Schematics of hetRTRs with interfaces colored corresponding to interface source next to nsEM class averages of (top left) A2B2-hetRTR05-11, (top right) A52B72-hetRTR05-11noCys including overlay of the 2D class with a 2D class of RTR11 to show angular shift, (bottom left) A3B3-hetRTR05-11noCys, and (bottom right) A2B2C2-hetRTR11noCys-05-16.

We tested 24 redesigns of RTR0, called RTR01 through RTR24, as homotetramers (Table S4). For 10/24 redesigns, SEC elution profiles suggested a single homogenous species with the target subunit composition (Fig. 5b, Fig. S13). Seven of those designs were primarily homotetramer according to nMS (Fig. S14; some monomer was also present in many of them), and for five of these SAXS profiles were consistent with the design model (Fig. 5c, Fig. S15). The interface cysteines of RTR11 and RTR24 were found not to be required for assembly (Fig. S15) so their cysteines were mutated to alanines in some later designs involving these interfaces to reduce the risk of disulfide-locked off-target species. Arm-like extensions protruding from the C-terminus of each subunit (Fig. 1d)^28^ could easily be identified by nsEM and facilitated determination of the oligomeric states (Fig. 5d, Fig. S16). The 2D class averages of RTR0 show a nearly perfectly circular ring with four studs spaced equally around the circle, corresponding to a top down view of the design model along its symmetry axis (Fig. S16). RTR05 and, to a lesser extent, RTR16 contained tetramers as well as pentamers, which had larger rings and five studs (Fig. 5d) and which were not apparent by nMS; no monomers or dimers were observed in any of these designs by nsEM. Any pseudosymmetric hetero-oligomers built using these homo-oligomers would be indistinguishable by nsEM, so to diversify the shape of the RTR we either extended the existing DHR extensions, attached DHR-DHR junctions with diverse shapes^31^ to the C-terminus, or used HFuse^32^ to attach miniproteins^33^ into the lumen on the N-terminus. We characterized these as extensions of RTR24 or RTR11 and found them to be clearly distinguishable (Fig. S17); by construction the extensions should be cross-compatible with all RTR variants.

We assessed the pairwise interface compatibility of the 5 redesigned homo-oligomers by building the full set of 10 C2-symmetric A2B2 heterotetramers (Table S5) via the same recombination protocol used for the BGLs, using Leu98 as the splicing junction. We designed the A2B2 heterotetramers so that each unique chain displayed a different C-terminal extension and were tagged with either GFP or a His6-tag. Many of the designs had multiple peaks on SEC (Fig. S18), but the peaks containing GFP (according to absorbance at 480 nm) in four of the designs, A2B2-hetRTR05-11, A2B2-hetRTR05-11_noCys_, A2B2-hetRTR05-18_noCys_, and A2B2-hetRTR18_noCys_-24_noCys_, were found to correspond to the intended A2B2 heterotetramers according to nMS (Fig. S19). This was confirmed for A2B2-hetRTR05-11 by nsEM, where the radial extensions indicate C2 symmetry (Fig. 5e upper left). We reasoned that the A2B2 complex should assemble successfully as long as the final assembly contained 24 total repeats in the ring, so to test this we modified the protomers to have one less or one more ring repeat (Chain A: 6 -> 5; Chain B:6->7;5×2 + 7×2 = 24). 3/6 tested designs (Table S5) still assembled (Fig. S18b), and nsEM of A_5_2B_7_2-hetRTR05-11_noCys_ revealed the studs shifted by approximately 15°, as expected for rotation by a single repeat (Fig. 5e top right)., A2B2 heterotetramers containing the RTR05 or RTR16 interfaces, which as homo-oligomers partially formed pentamers, did not contain any off-target pentameric species, suggesting that the added specificity constraints of heteromeric interactions may prevent off-target assemblies. This has also been observed in heteromeric assemblies of native LSm proteins, which have reduced variation in ring size compared to related homomeric assemblies.^5^

To explore the creation of oligomers with different numbers of subunits, we designed homomeric C3, C6, C8, and C12 versions of the RTR toroid (Table S4). The C3 versions assembled as desired, but complexes with C6 symmetry and above assembled into a range of oligomers with different numbers of subunits (Fig. S12). As noted above, we hypothesized that the heteromeric versions of these assemblies might better assemble into the target stoichiometries than their homomeric counterparts. Therefore, we created heterohexameric rings using the recombination approach which combined either two different interfaces into A3B3 hexamers, or three interfaces into A2B2C2 hexamers, employing the interfaces found to be successful in the A2B2 designs. The same hybrid protomers could be used in both A3B3 or A2B2C2 hexamers as the combination of interfaces programs the final assembly. When expressed separately and mixed (Table S6), 3/12 tested A3B3 combinations assembled, and 2/4 tested A2B2C2 combinations assembled, as measured by SEC (Fig. S20). A3B3-hetRTR05-11_noCys_ was verified by nMS (Fig. S19), and the expected ring shape with two different alternating extensions was observed by nsEM (Fig. 5e, lower left). We were also able to confirm one A2B2C2 assembly by nsEM where three different extensions with the intended circular arrangement can be observed (Fig. 5e, bottom right). In both cases, the heterohexameric designs were found to be homogenous in their oligomeric state, in contrast with the polydispersity found in their homohexameric parental designs.

## Discussion

*De novo* hetero-oligomers, especially with three or more unique chains, are powerful building blocks for enabling the design of larger and more sophisticated nanostructures and devices.^18^ They are, however, challenging to create because they require the design of multiple interfaces that all need to assemble correctly within one oligomer; failure of one interface leads to complete failure of the hetero-oligomer even if the other interface form correctly. This compounded probability of failure generally means testing many, sometimes hundreds, of designs to find a few which assemble properly.^11,16^ By first evaluating each interface separately in a homo-oligomeric context and then generating hetero-oligomers by structural splicing, our hierarchical homologous recombination-inspired design strategy yielded 17 experimentally-verified hetero-oligomers (out of 49 tested designs). This success was achieved by first building and validating a small set of homo-oligomers to draw from. The BGL family was particularly amenable to interface redesign, and we were able to generate 10 new robust and specific ABC heterotrimers (of 17 tested) which considerably expand the possibilities for pseudosymmetric nanomaterials (see below). The RTR family yielded 4 (of 10 tested) A2B2s, of which the combination of interfaces from RTR05 and RTR11_noCys_ was particularly robust and were used as the basis for the successful A3B3 (1 of 12 tested) and A2B2C2 (1 of 4 tested) designs. The structural versatility of the RTR family arising from the ability to independently and modularly change the stoichiometries, chain identities, shapes, and spacings of their DHR extensions/termini provides a range of diverse hubs for target clustering in cell signaling and other applications^9,34^. The progenitor of this family was used as a component of designed molecular machines^28^, and the pseudosymmetric RTRs could be readily incorporated into more complex molecular machines. In this work, we applied our pseudosymmetrization strategy to *de novo* designed assemblies; it would be straightforward to generalize the approach to natural homomeric proteins of interest, such as TNF ligands which might induce novel signaling responses^35^, or to create hetero-oligomers with four or more unique subunits.

Pseudosymmetry provides a means to easily transition between simple homo-oligomeric systems and more complicated hetero-oligomeric systems. Fusions or other modifications to hetero-oligomers can first be prototyped using their homo-oligomeric paralogs which are often easier to validate, as demonstrated by the diversification of the RTR extensions by fusion of DHR junctions (Fig. S17). Homo-oligomers with pseudosymmetric variants can similarly be used for material design by initially building higher order symmetric assemblies out of homo-oligomers and, as a second step, breaking symmetry by substituting the homo-oligomers for their pseudosymmetric hetero-oligomeric variants, which enables the specialization of a subset of the interactions while preserving the interaction geometry between the pseudosymmetric building blocks. This opens up routes to more complex protein nanomaterials; for example, as described in the accompanying preprint, we used the BGL homotrimers to design simple tetrahedral, octahedral, and icosahedral nanocages which we then transformed into more complex T=4 nanocages by substituting in hetBGL trimers to desymmetrize the nanocage, preserving the interface geometries of two of the interfaces and modifying the third to interact with a new homotrimer which increases the triangulation number, physical size, and complexity of the nanocage^1^.

## Acknowledgements

We would like to thank Nick Woodall, Phil Bradley, Scott Boyken, and Zibo Chen for helpful discussions, Andrew Borst for EM support, and Kandise Van Wormer and Luki Goldschmidt for technical support. We also thank Drs. Rosa Viner and Shane Bechler (Thermo Fisher Scientific, Sunnyvale, CA) for providing the desalting cartridge prototypes used in this study.

Native mass spectrometry measurements were provided by Florian Busch, Andrew Norris, Nicholas Horvath, Stella Lai, Sean Cleary, and Marius Kostelic of the NIH-funded Resource for Native Mass Spectrometry Guided Structural Biology at The Ohio State University (NIH P41 GM128577 awarded to Vicki Wysocki).

SAXS data collection was provided by Kathryn Burnett and Greg Hura via the SIBYLS mail-in SAXS program at the Advanced Light Source (ALS), a national user facility operated by Lawrence Berkeley National Laboratory on behalf of the Department of Energy, Office of Basic Energy Sciences, through the Integrated Diffraction Analysis Technologies program, supported by the Department of Energy (DOE) Office of Biological and Environmental Research. Additional support comes from the NIH project ALS-ENABLE (grant P30 GM124169) and High-End Instrumentation Grant S10OD018483.

X-ray crystallography diffraction data were collected on a Pilatus area detector at the Advanced Light Source (ALS) synchrotron facility at beamline 5.0.2 for BGL06, BGL14_styr, BGL15, and BGL18. Diffraction data were collected on a Rigaku HyPix-6000HE hybrid photon counting detector at the Fred Hutchinson Cancer Center (FHCC) for hetBGL03-15-18.

## Funding sources

The Audacious Project (RDK, LC)

The Nordstrom Barrier Institute for Protein Design Directors Fund (CMC)

Bill and Melinda Gates Foundation #OPP1156262 (LC, XL)

SSGCID is supported by National Institute of Allergy and Infectious Diseases (NIAID)

Federal Contract # HHSN272201700059C (LC)

The Open Philanthropy Project Improving Protein Design Fund (BIMW, LC, CMC)

EMBO long-term fellowship (ALTF 139-2018) (BIMW)

NSFGRFP DGE-1762114 (MAK)

NIH R01 GM105691 (BLS)

NIH shared instrumentation grant S10OD021832 (BLS)

NIH P41 GM128577 (VHW, SML, MMK)

## Competing interests

A provisional patent application has been filed (63/486,872) by the University of Washington, listing R.D.K., Nicholas Woodall, B.I.M.W., B.L.S., S.L., and D.B. as inventors or contributors.

