## Supplemental Information for "Stepwise design of pseudosymmetric protein hetero-oligomers"

### Supplemental Data

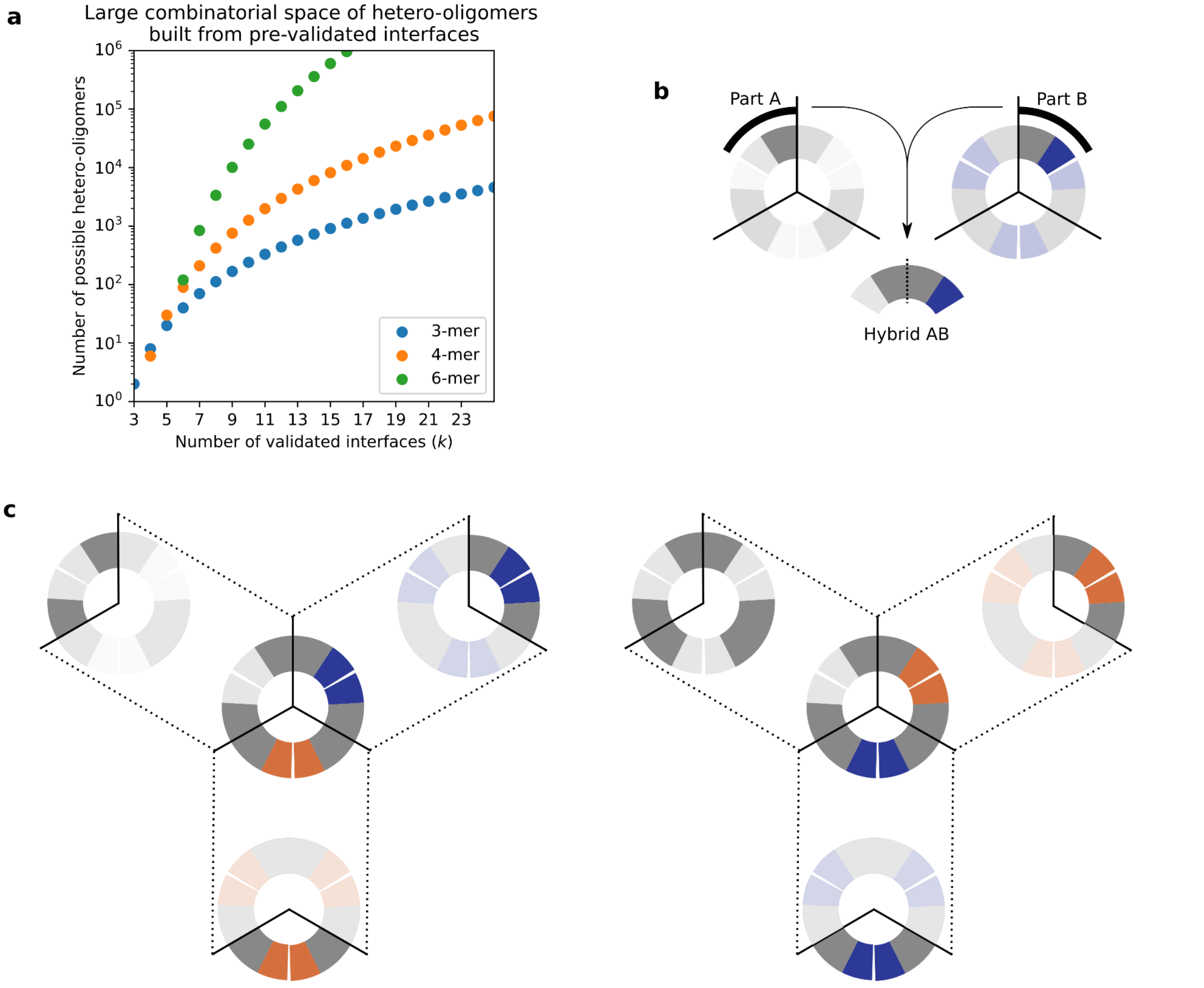


**Supplemental Figure 1: Combinatorics of homo-oligomer recombination**

(**a**) Plot of the equation describing the number of possible hetero-oligomers that can be made, $(k! * (n-1)!) / (n! * (k-n)!)$, for a few n, where n is the number of different chains in the final heter-n-mer (e.g. heterotrimer = heter-3-mer) and k is the number of interfaces or homo-n-mers which have been validated to assemble similarly. Shown are plots of n=3, 4, or 6, some of the most commonly useful hetero-oligomers, to illustrate the large number of options for combining a few interfaces. (b) The method of combining parts of two homotrimers to form a hybrid protomer of a heterotrimer. The chains of the homotrimers are computationally split into two parts at the same point on each chain as shown by the 3-pointed crosshair overlays, creating part A and part B. Because the molecular context at the cut points are identical, parts A and B can be spliced together at the cut point to form Hybrid AB without altering the structure of the protomer. (c) There are two ways to combine the same three homotrimers to form pseudosymmetric heterotrimers because order matters for the way the interfaces are arranged around the circle. There are many more ways to combine higher n hetero-n-mers.

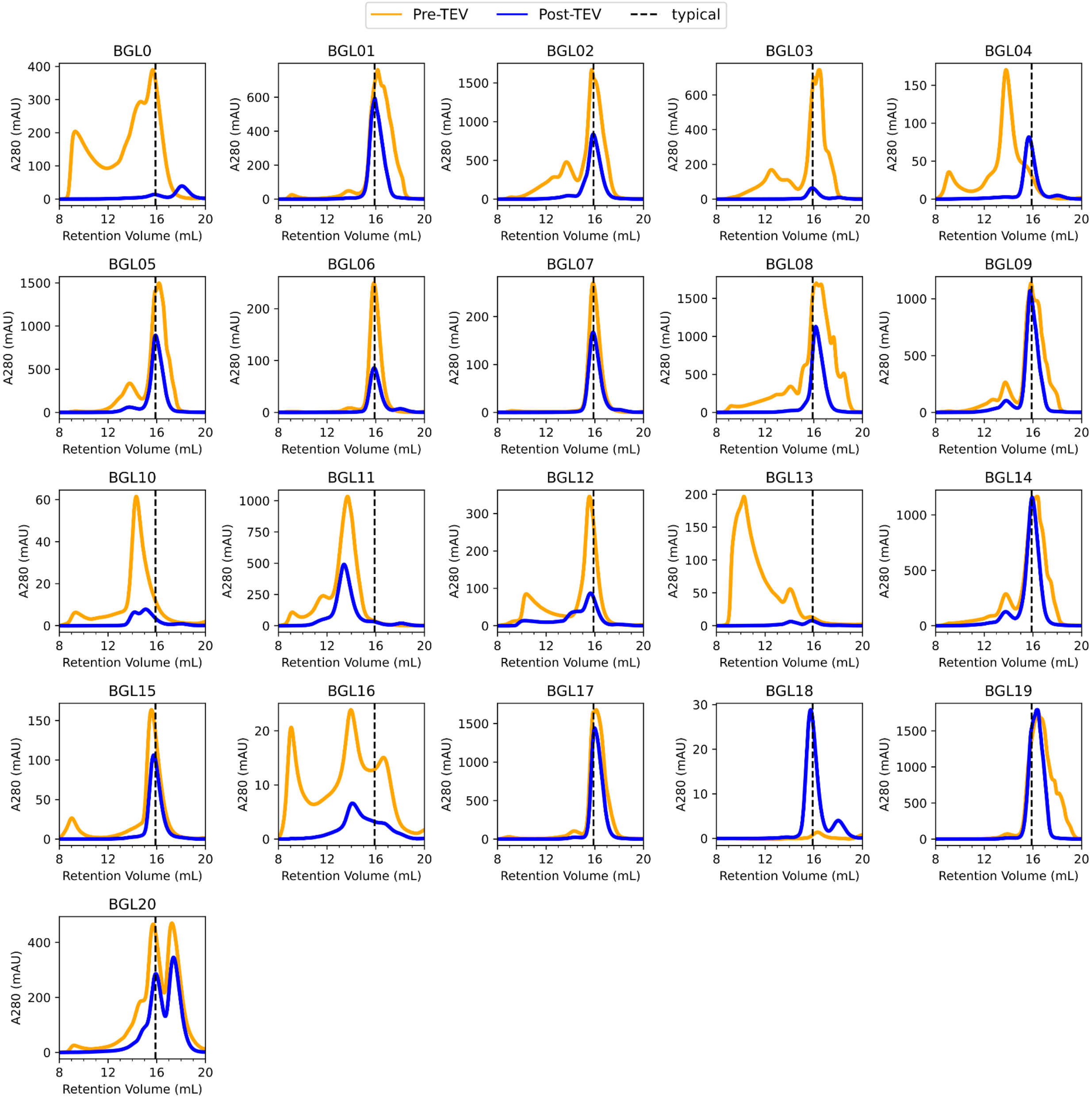

**Supplemental Figure 2: SEC of BGL homotrimers**

Plots of all designs before (orange) and after (blue) TEV cleavage, run on S200 Increase 10/300 GL. Note that some samples were split and run separately so absolute abundance is not comparable in all cases. The right-shift pre- to post-TEV in BGL04 can be explained by removal of interactions via mediated His6-tags, which are a relatively common occurrence in cyclic homo-oligomers. BGL0 was not efficiently cleaved, causing low yield of possibly mis-assembled protein in post-TEV SEC.

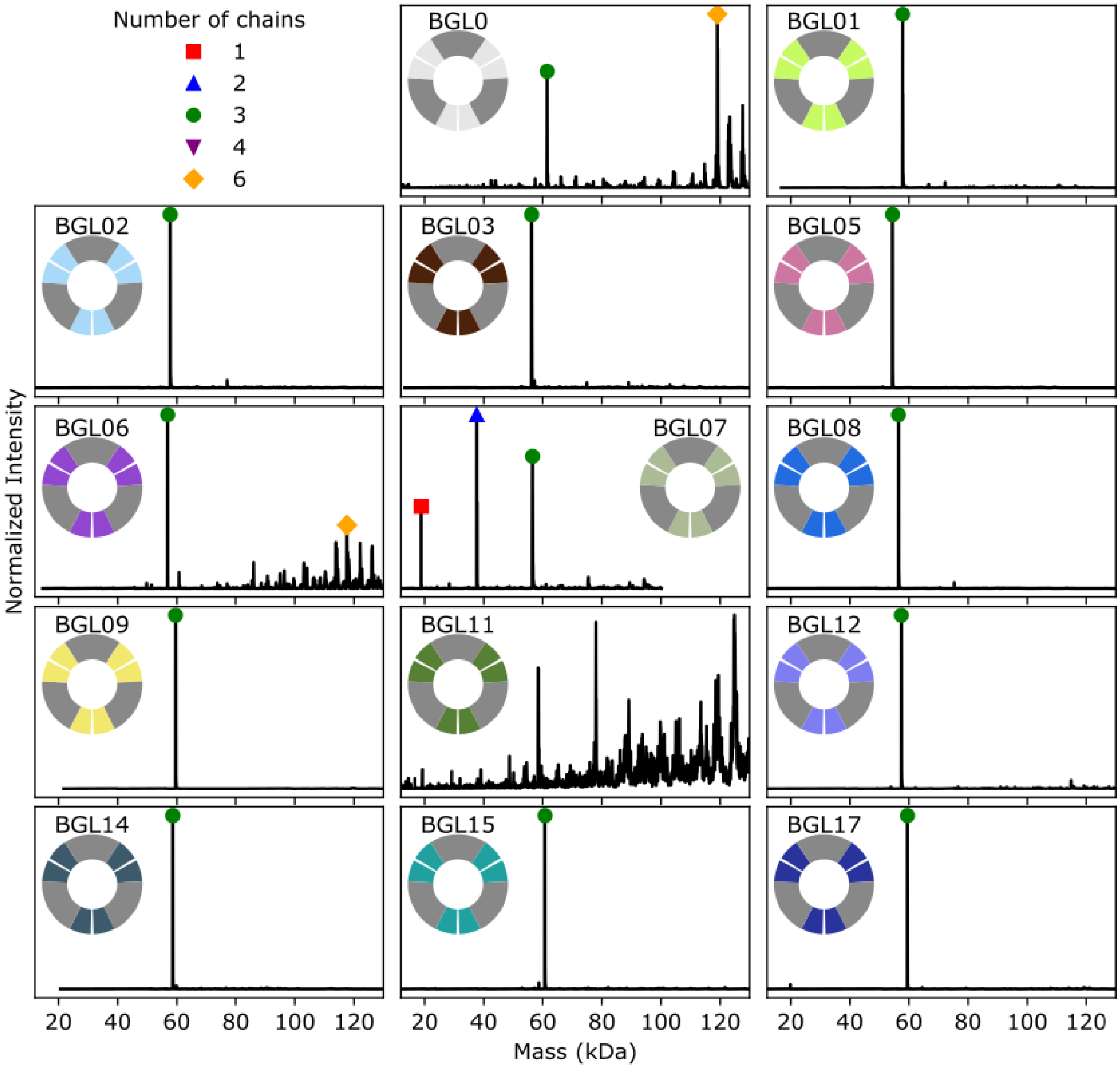

**Supplemental Figure 3: nMS of BGL homotrimers**

Plots of all designs with nMS data. Black, deconvoluted masses obtained from m/z spectra (see Supplemental Table 7). Identified species corresponding to monomer (red square), dimer (blue triangle), trimer (green circle), tetramer (purple upside-down triangle), and hexamer (yellow diamond) are annotated; other un-annotated peaks are unidentified species.

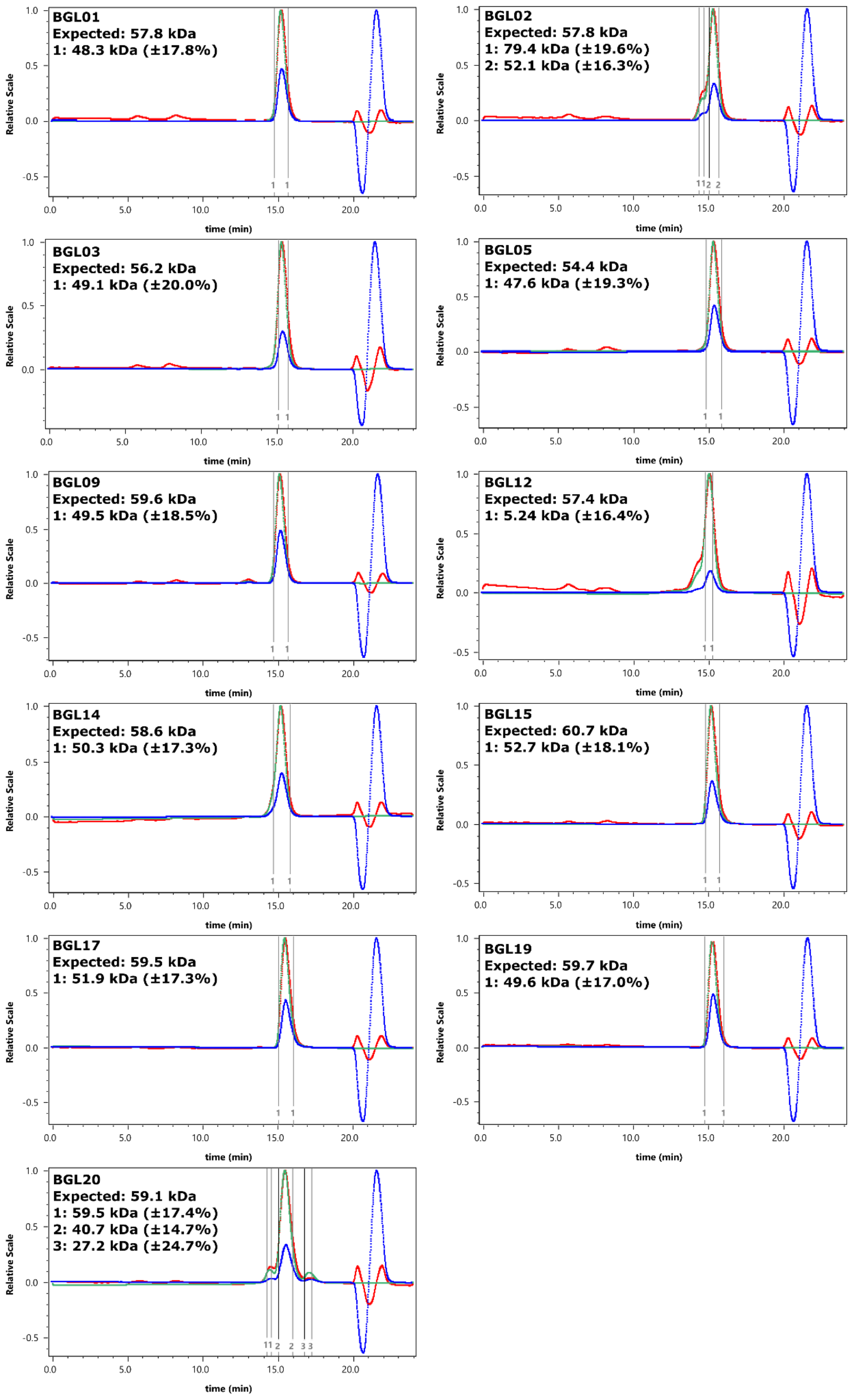


**Supplemental Figure 4: SEC-MALS of homotrimers**

Plots of all designs with SEC-MALS data. Red, light scattering signal. Green, UV absorbance. Blue, differential refractive index (dRI). Peaks are labeled by goalposts. Molecular weight estimates for each peak are shown in the top right corner of each plot.

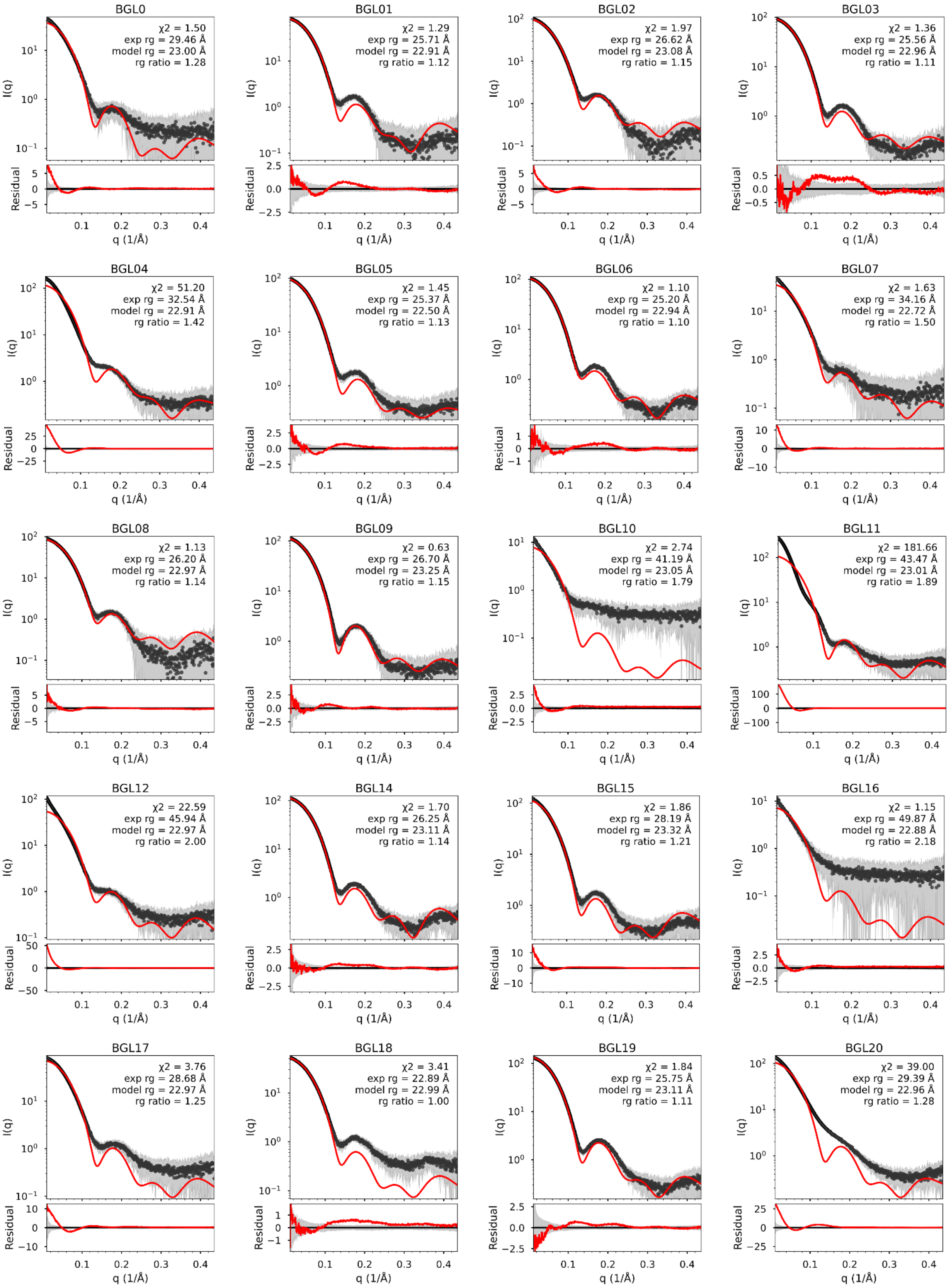
BGL0
χ2 = 1.50
exp rg = 29.46 Å
model rg = 23.00 Å
rg ratio = 1.28
BGL01
χ2 = 1.29
exp rg = 25.71 Å
model rg = 22.91 Å
rg ratio = 1.12
BGL02
χ2 = 1.97
exp rg = 26.62 Å
model rg = 23.08 Å
rg ratio = 1.15
BGL03
χ2 = 1.36
exp rg = 25.56 Å
model rg = 22.96 Å
rg ratio = 1.11
BGL04
χ2 = 51.20
exp rg = 32.54 Å
model rg = 22.91 Å
rg ratio = 1.42
BGL05
χ2 = 1.45
exp rg = 25.37 Å
model rg = 22.50 Å
rg ratio = 1.13
BGL06
χ2 = 1.10
exp rg = 25.20 Å
model rg = 22.94 Å
rg ratio = 1.10
BGL07
χ2 = 1.63
exp rg = 34.16 Å
model rg = 22.72 Å
rg ratio = 1.50
BGL08
χ2 = 1.13
exp rg = 26.20 Å
model rg = 22.97 Å
rg ratio = 1.14
BGL09
χ2 = 0.63
exp rg = 26.70 Å
model rg = 23.25 Å
rg ratio = 1.15
BGL10
χ2 = 2.74
exp rg = 41.19 Å
model rg = 23.05 Å
rg ratio = 1.79
BGL11
χ2 = 181.66
exp rg = 43.47 Å
model rg = 23.01 Å
rg ratio = 1.89
BGL12
χ2 = 22.59
exp rg = 45.94 Å
model rg = 22.97 Å
rg ratio = 2.00
BGL14
χ2 = 1.70
exp rg = 26.25 Å
model rg = 23.11 Å
rg ratio = 1.14
BGL15
χ2 = 1.86
exp rg = 28.19 Å
model rg = 23.32 Å
rg ratio = 1.21
BGL16
χ2 = 1.15
exp rg = 49.87 Å
model rg = 22.88 Å
rg ratio = 2.18
BGL17
χ2 = 3.76
exp rg = 28.68 Å
model rg = 22.97 Å
rg ratio = 1.25
BGL18
χ2 = 3.41
exp rg = 22.89 Å
model rg = 22.99 Å
rg ratio = 1.00
BGL19
χ2 = 1.84
exp rg = 25.75 Å
model rg = 23.11 Å
rg ratio = 1.11
BGL20
χ2 = 39.00
exp rg = 29.39 Å
model rg = 22.96 Å
rg ratio = 1.28
I(q)
Residual
q (1/Å)

**Supplemental Figure 5: SAXS profiles of BGL homotrimers**

Plots of all designs with SAXS data. (Top of each panel) The scattering profiles plotting log intensity vs scattering angle. Black dots, averaged experimental data from many individual frames. Gray shading, the standard deviation of the averaged frames. Red, computed profiles generated from design models and scaled using FoXS to fit the experimental data with c1=1.05 and c2=2.0. (Bottom of each panel) The residual of the fit between experimental data and the computed profile, shown on a linear scale. Red, magnitude of the residual. Gray shading, same as top. Quality of fit ($X^2$), radius of gyration (rg) from experimental data and the design model, and the ratio between the two, are listed in the top right of each panel.

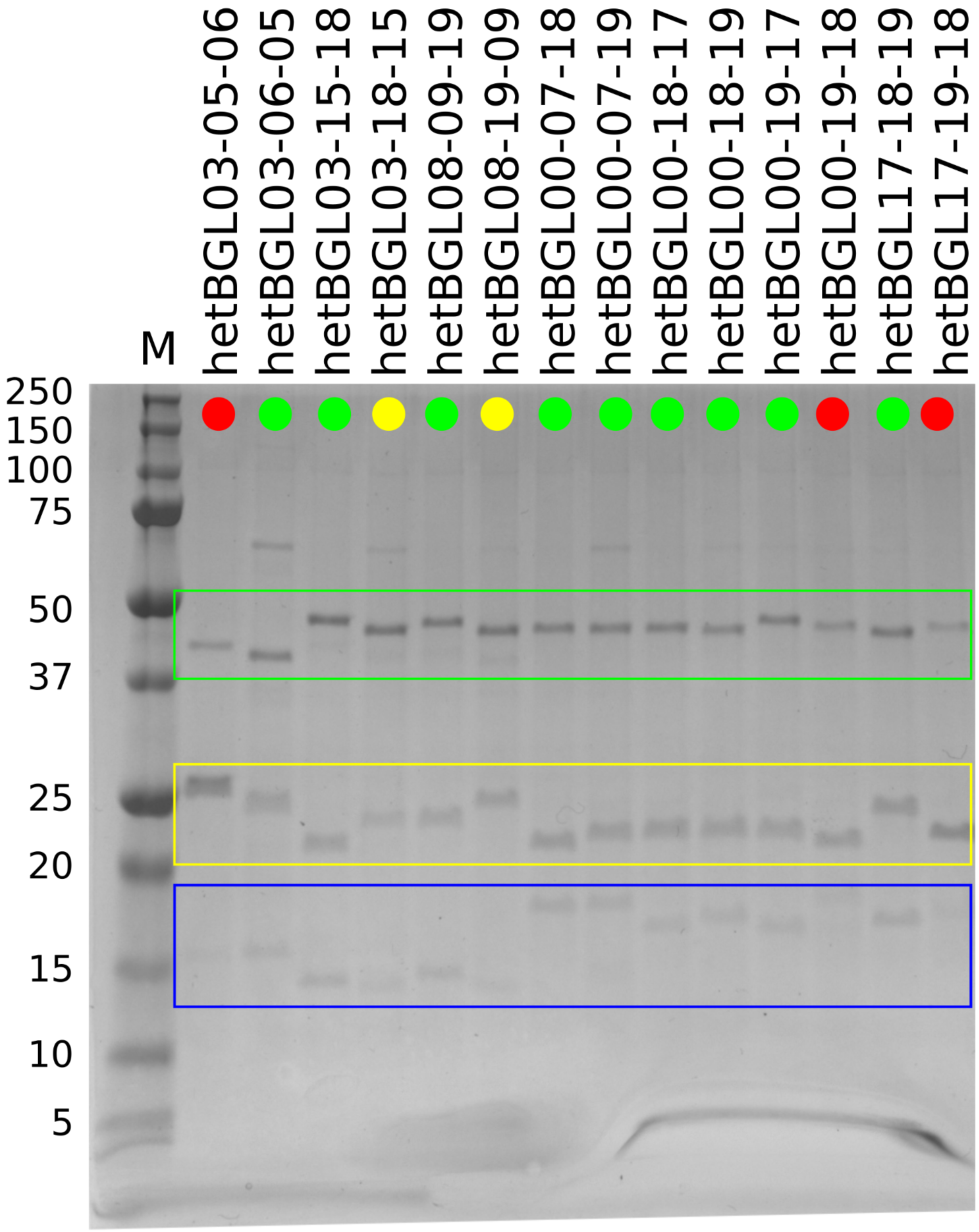


**Supplemental Figure 6: SDS-PAGE of hetBGL co-purifications**

IMAC co-purified co-expressions, where each construct only contains a single His6-tag. Refer to Supplemental Table 3 for His6-tag placement. The three chains are fused to different mass-adding tags to differentiate them. Green box, GFP-tagged. Yellow box, EHEE-tagged. Blue box, no mass tag. M lane is Precision Plus Protein Dual Xtra (Bio-Rad) and listed masses are in kDa. Sample lanes are annotated with dots to indicate qualitative success. Red dot, missing chain. Yellow dot, all 3 chains present but uneven stoichiometries. Green dot, all 3 chains present in approximately equal stoichiometries.

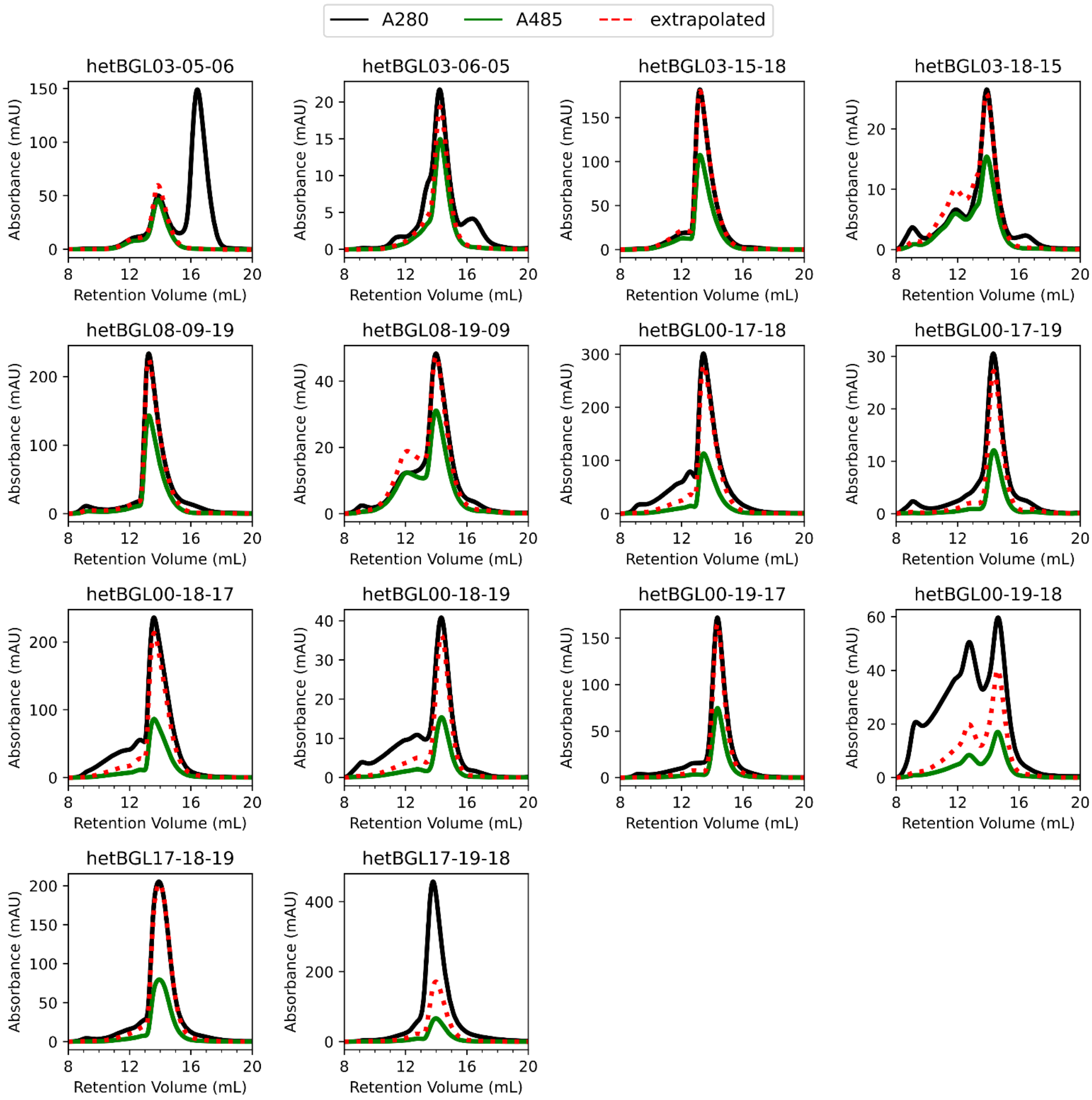
A280
A485
extrapolated
hetBGL03-05-06
hetBGL03-06-05
hetBGL03-15-18
hetBGL03-18-15
hetBGL08-09-19
hetBGL08-19-09
hetBGL00-17-18
hetBGL00-17-19
hetBGL00-18-17
hetBGL00-18-19
hetBGL00-19-17
hetBGL00-19-18
hetBGL17-18-19
hetBGL17-19-18
Absorbance (mAU)
Retention Volume (mL)

**Supplemental Figure 7: SEC of hetBGLs**

Each successfully assembling construct will have a single GFP tag associated with it, so tracking A280 (absorbance of aromatics in every chain) and A485 (absorbance specific to the GFP chromophore) separately allows for estimation of expected signal given another signal which implies GFP:complex stoichiometry. Black, measured A280. Green, measured A485. Red dashed, expected A280 signal given the A485 signal using an estimated EC485 of 36650 $M^{-1}cm^{-1}$ and estimated A280 specific to each construct.

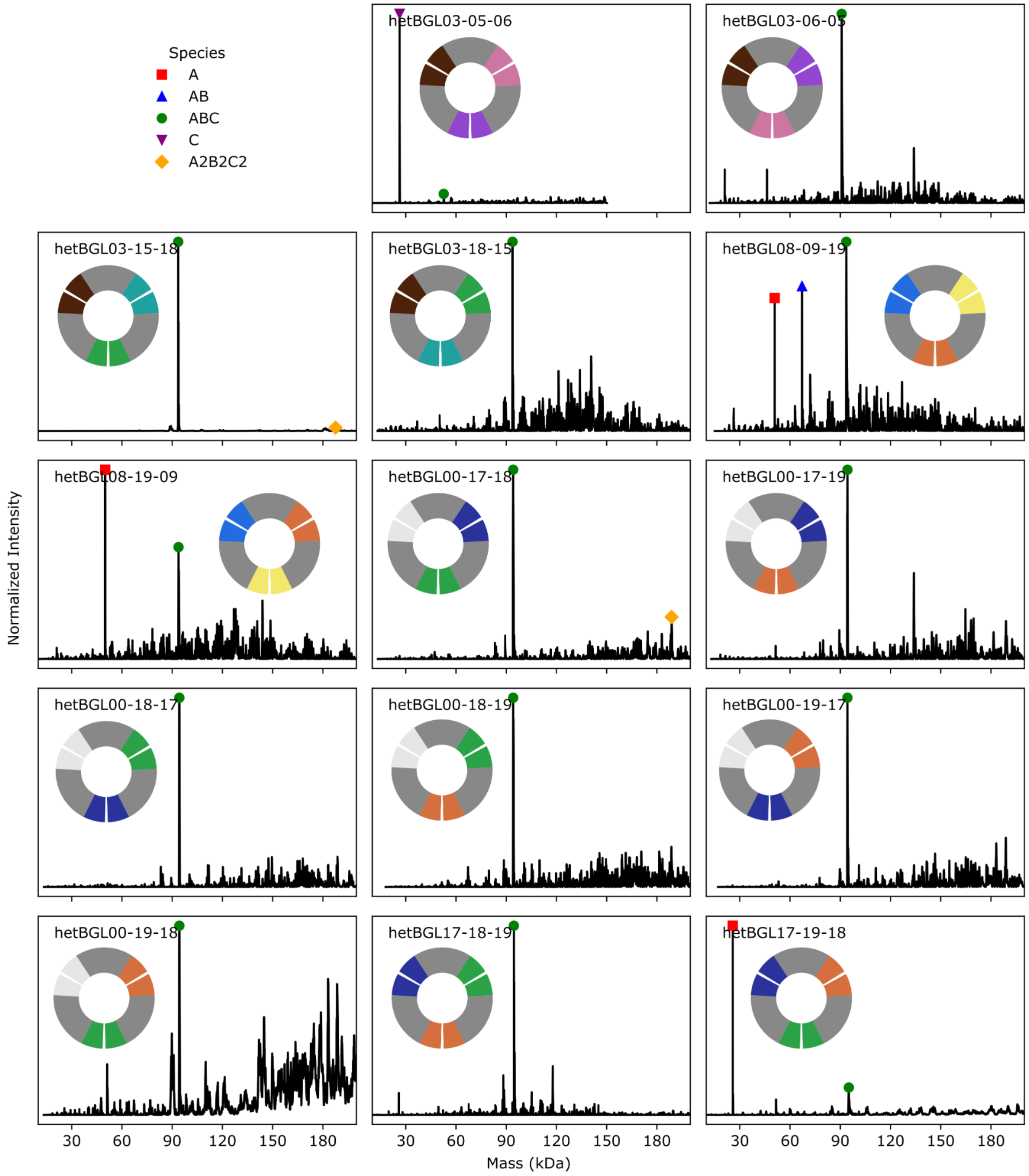


**Supplemental Figure 8: nMS of hetBGLs**

Black, deconvoluted masses obtained from m/z spectra (see Supplemental Table 7). Identified species corresponding to chain A (red square), A+B (blue triangle), A+B+C (green circle), C (purple upside-down triangle), and (A+B+C)x2 (yellow diamond) are annotated; other un-annotated peaks are unidentified species.

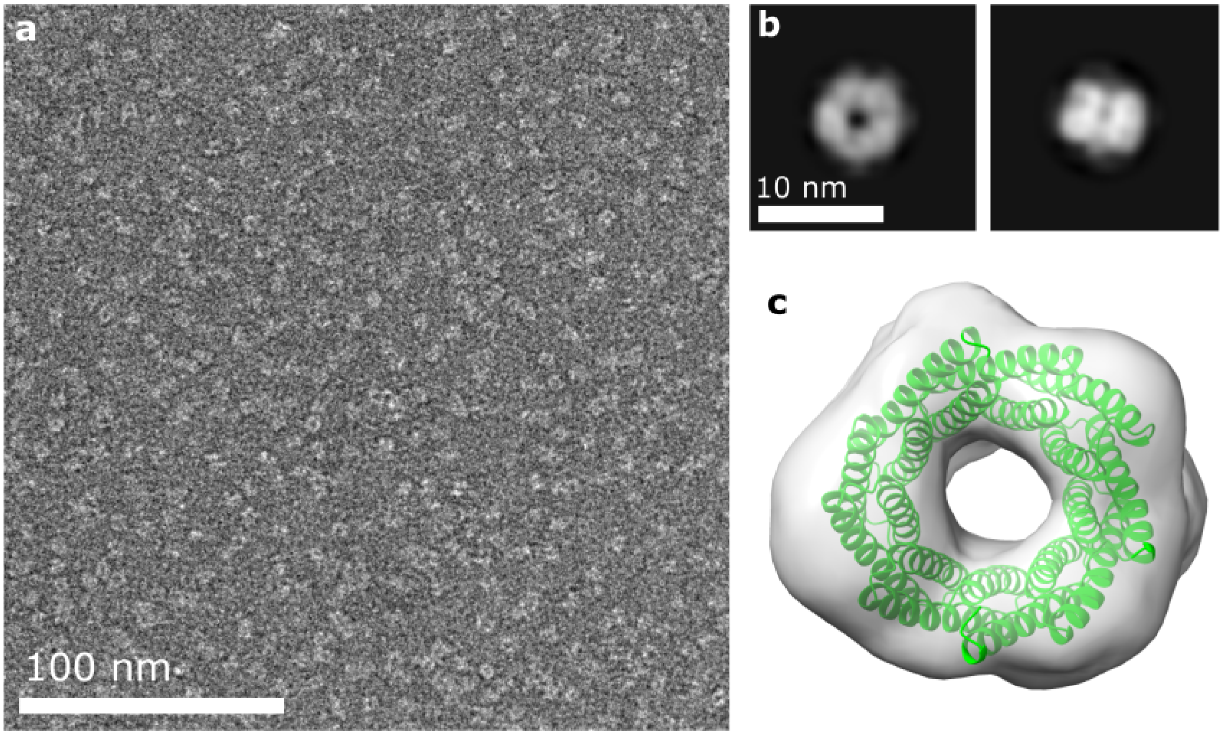


**Supplemental Figure 9: nsEM of hetBGL03-15-18**

(a) Randomly chosen example micrograph. (b) Two examples 2D class averages showing the top-down view and a probable side view. (c) Design model fit into 3D reconstruction.

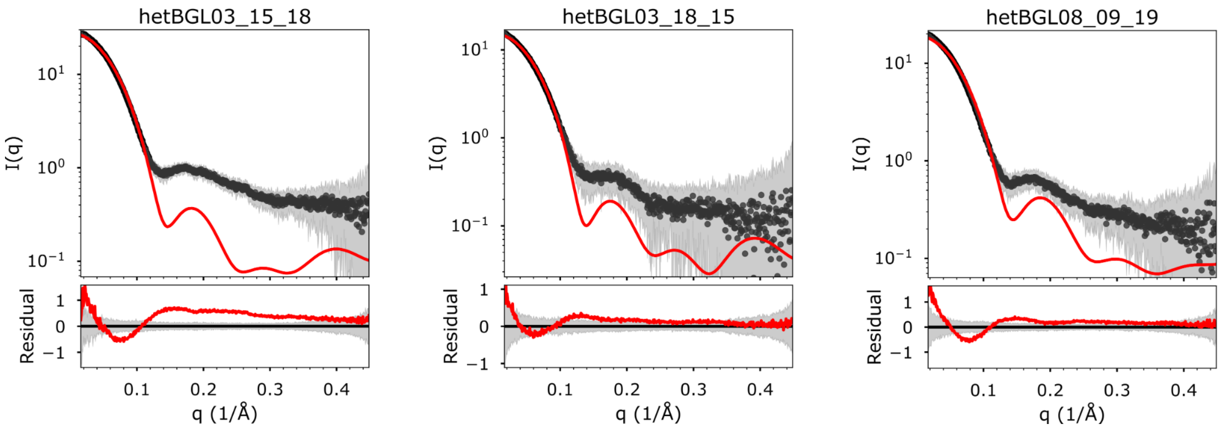


**Supplemental Figure 10: SAXS of hetBGLs**

Plots of all designs with SAXS data. (Top of each panel) The scattering profiles plotting log intensity vs scattering angle. Black dots, averaged experimental data from many individual frames. Gray shading, the standard deviation of the averaged frames. Red, computed profiles generated from design models and scaled using FoXS to fit the experimental data. (Bottom of each panel) The residual of the fit between experimental data and the computed profile, shown on a linear scale. Red, magnitude of the residual. Gray shading, same as top. Quality of fit ($X^2$), radius of gyration (rg) from experimental data and the design model, and the ratio between the two, are listed in the top right of each panel.

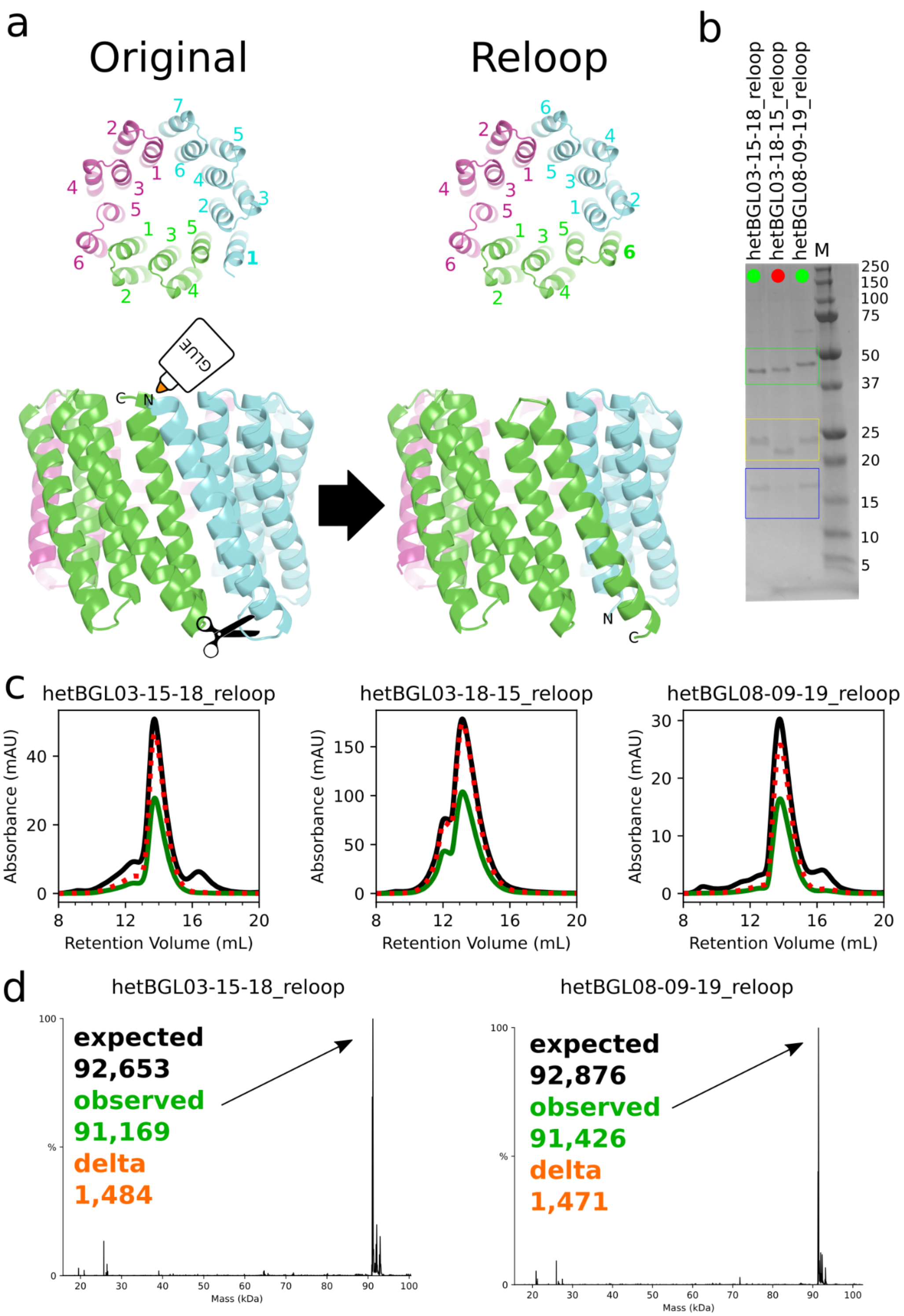


**Supplemental Figure 11: Circular permutation of Set 1 hetBGLs**

(a) Set 1 hetBGLs ("Original") have different numbers of helices in each chain. To convert them into structures more resembling those of Set 2 which have six helices per chain, the loop between the first and second helix of the longest chain (blue, left) was cut and attached to the

C-terminus of the shortest chain (green) using the same loop design procedure used to make the HR-C homotrimers. The result is the heterotrimer on the right with six helices per chain and where all chains have the same topology. (b) SDS-PAGE of IMAC co-purified co-expressions of relooped hetBGLs, where each construct only contains a single His6-tag. Refer to Supplemental Table 3 for His6-tag placement. The three chains are fused to different mass-adding tags to differentiate them. Green box, GFP-tagged. Yellow box, EHEE-tagged. Blue box, no mass tag. M lanes are Precision Plus Protein Dual Xtra (Bio-Rad) and listed masses are in kDa. Sample lanes are annotated with dots to indicate qualitative success. Red dot, missing chain. Yellow dot, all 3 chains present but uneven stoichiometries. Green dot, all 3 chains present in approximately equal stoichiometries. (c) SEC traces. Black, measured A280. Green, measured A485. Red dashed, expected A280 signal given the A485 signal using an estimated EC485 of 36650 $M^{-1}cm^{-1}$ and estimated A280 specific to each construct. (d) Black, deconvoluted masses obtained from m/z spectra (see Supplemental Table 7). The major peaks are indicated with arrows with masses listed in the top left corner of each plot.

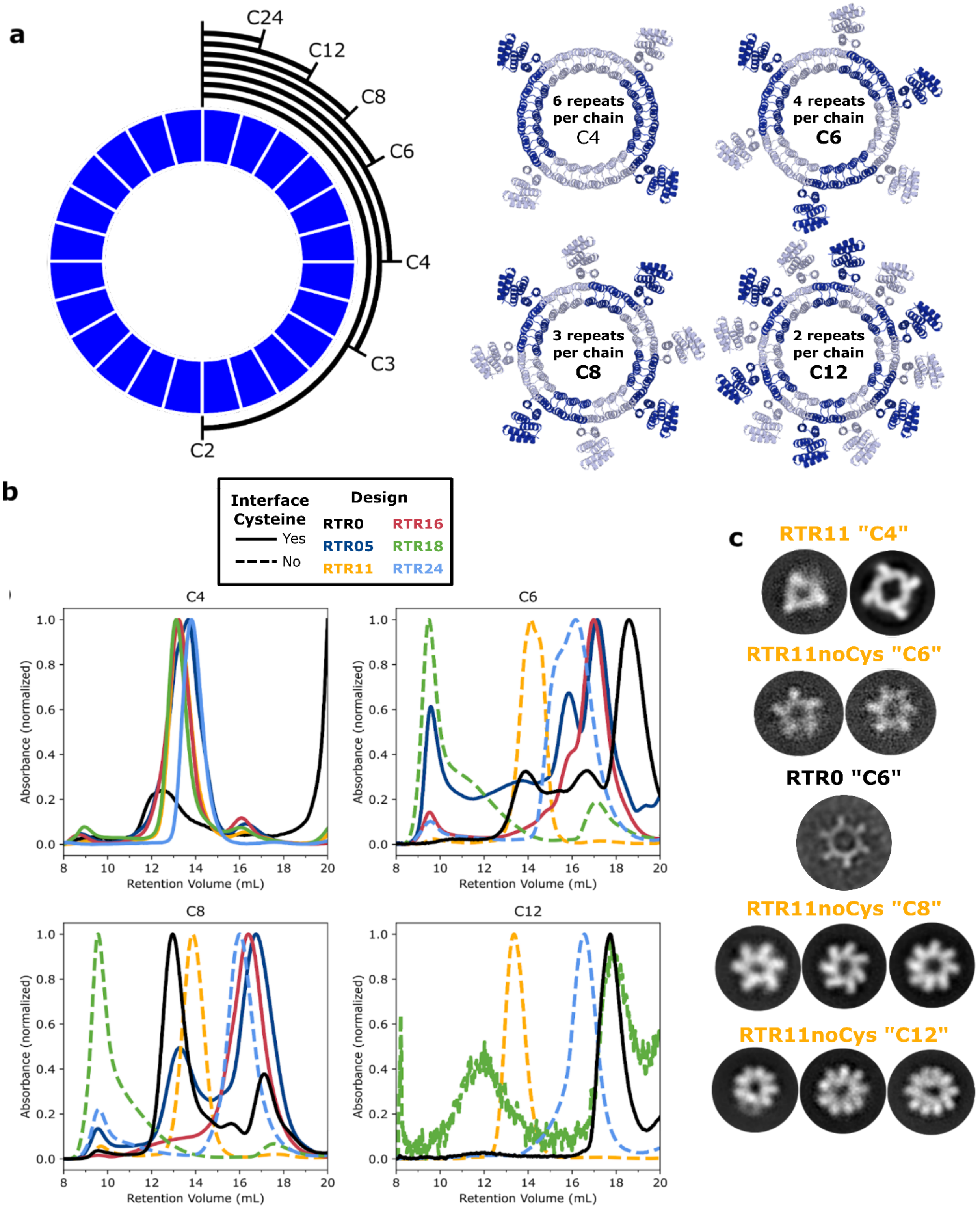

a
C24
C12
C8
C6
C4
C3
C2
6 repeats per chain C4
4 repeats per chain C6
3 repeats per chain C8
2 repeats per chain C12
b
Interface Cysteine
Yes
No
Design
RTR0
RTR16
RTR05
RTR18
RTR11
RTR24
C4
C6
C8
C12
Absorbance (normalized)
Retention Volume (mL)
c
RTR11 "C4"
RTR11noCys "C6"
RTR0 "C6"
RTR11noCys "C8"
RTR11noCys "C12"

**Supplemental Figure 12: Variable oligomeric state of RTR homo-oligomers**

(a, left) Schematic representation of how a cyclic (24x) repeat protein (toroid) can be divided into different homo-oligomers which assemble into symmetries depending on the number of repeats combined into a single chain. The final assemblies all have the same number of total repeats. For example, a C4 assembly is made from a single chain containing 6 repeats. (right) Cartoon models of C4, C6, C8, and C12 versions of the RTR backbone, constructed by removing repeats distant from the interface repeats. (b) SEC traces of RTR variants with different numbers of repeats per chain. Solid traces, interfaces with cysteines. Dashed traces, interfaces without cysteines. Black, RTR0. Dark blue, RTR05. Yellow, RTR11. Red, RTR16. Green, RTR18. Light blue, RTR24. (c) Negative stain EM 2D class averages of RTR0 or RTR11 with different numbers of repeats per chain. Each class represents a different observed oligomeric state.

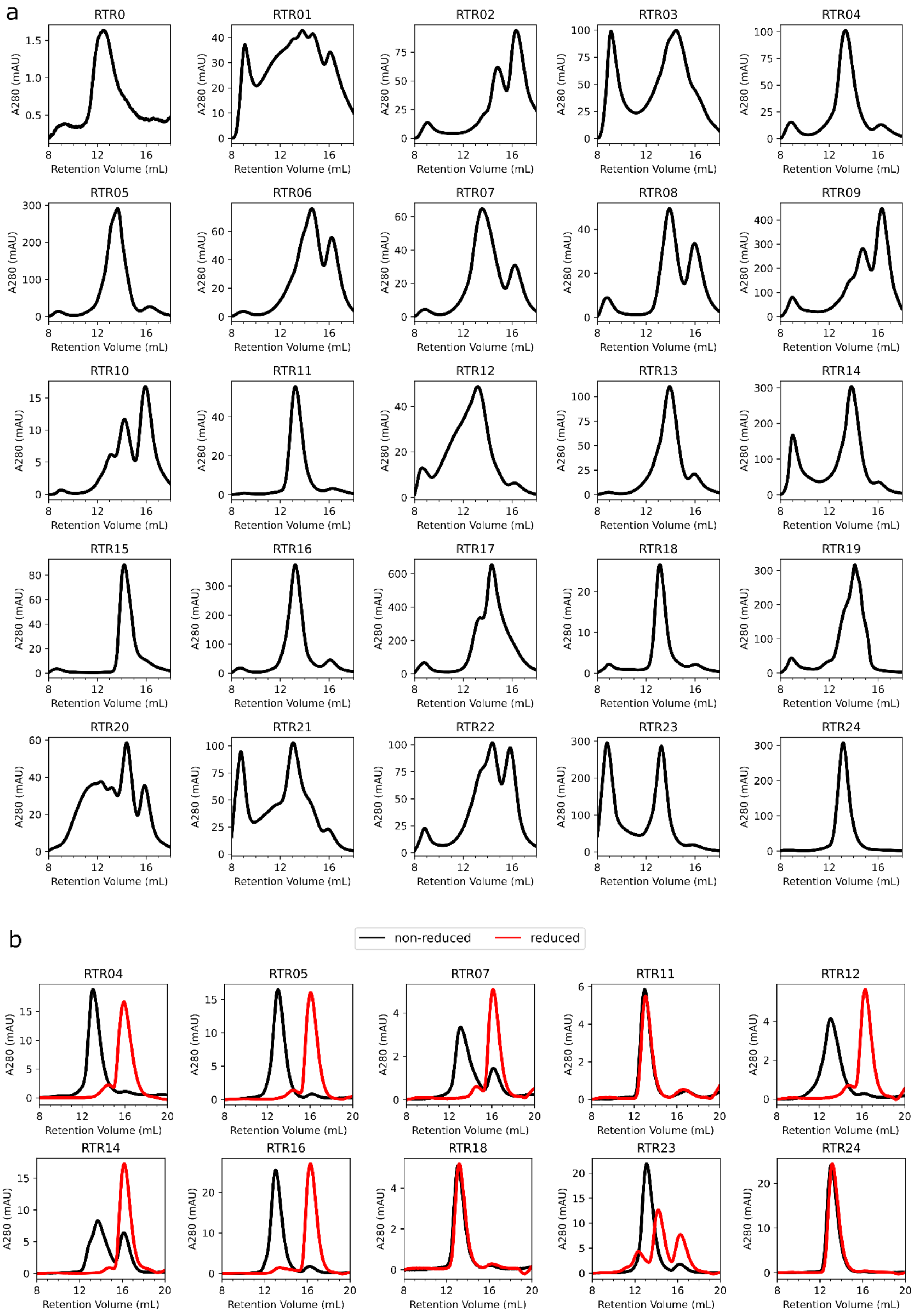
a
RTR0
RTR01
RTR02
RTR03
RTR04
RTR05
RTR06
RTR07
RTR08
RTR09
RTR10
RTR11
RTR12
RTR13
RTR14
RTR15
RTR16
RTR17
RTR18
RTR19
RTR20
RTR21
RTR22
RTR23
RTR24
A280 (mAU)
Retention Volume (mL)
b
non-reduced
reduced
RTR04
RTR05
RTR07
RTR11
RTR12
RTR14
RTR16
RTR18
RTR23
RTR24
A280 (mAU)
Retention Volume (mL)

**Supplemental Figure 13: SEC of RTR homo-tetramers**

SEC traces of (a) initial RTR samples on S200 column and (b) reran isolated fractions after being reduced with 10mM TCEP or not.

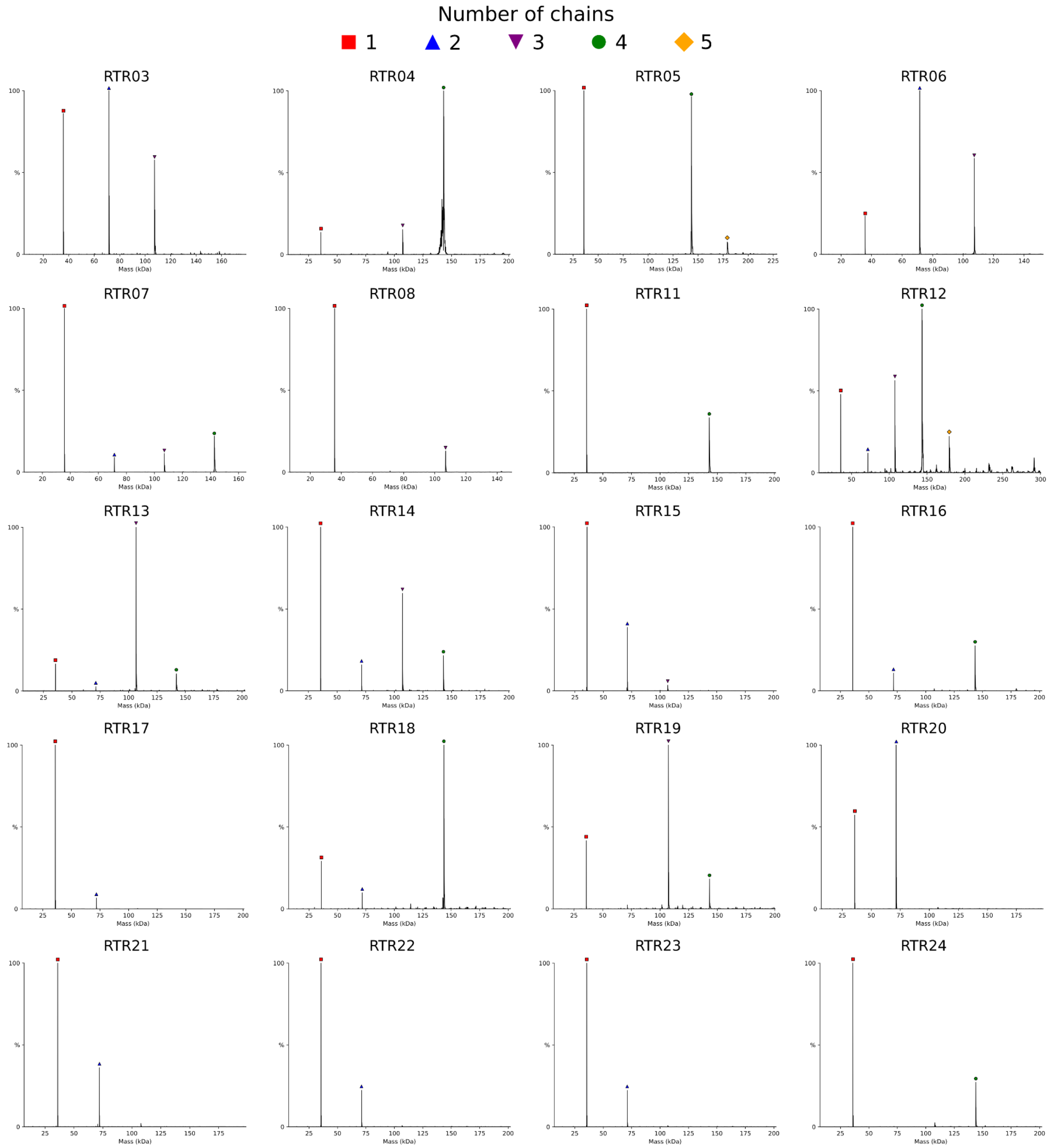


**Supplemental Figure 14: nMS of RTR homo-tetramers**

Plots of all designs with nMS data. Black, deconvoluted masses obtained from m/z spectra (see Supplemental Table 7). Identified species corresponding to monomer (red square), dimer (blue triangle), trimer (purple upside-down triangle), tetramer (green circle), and hexamer (yellow diamond) are annotated; other un-annotated peaks are unidentified species.

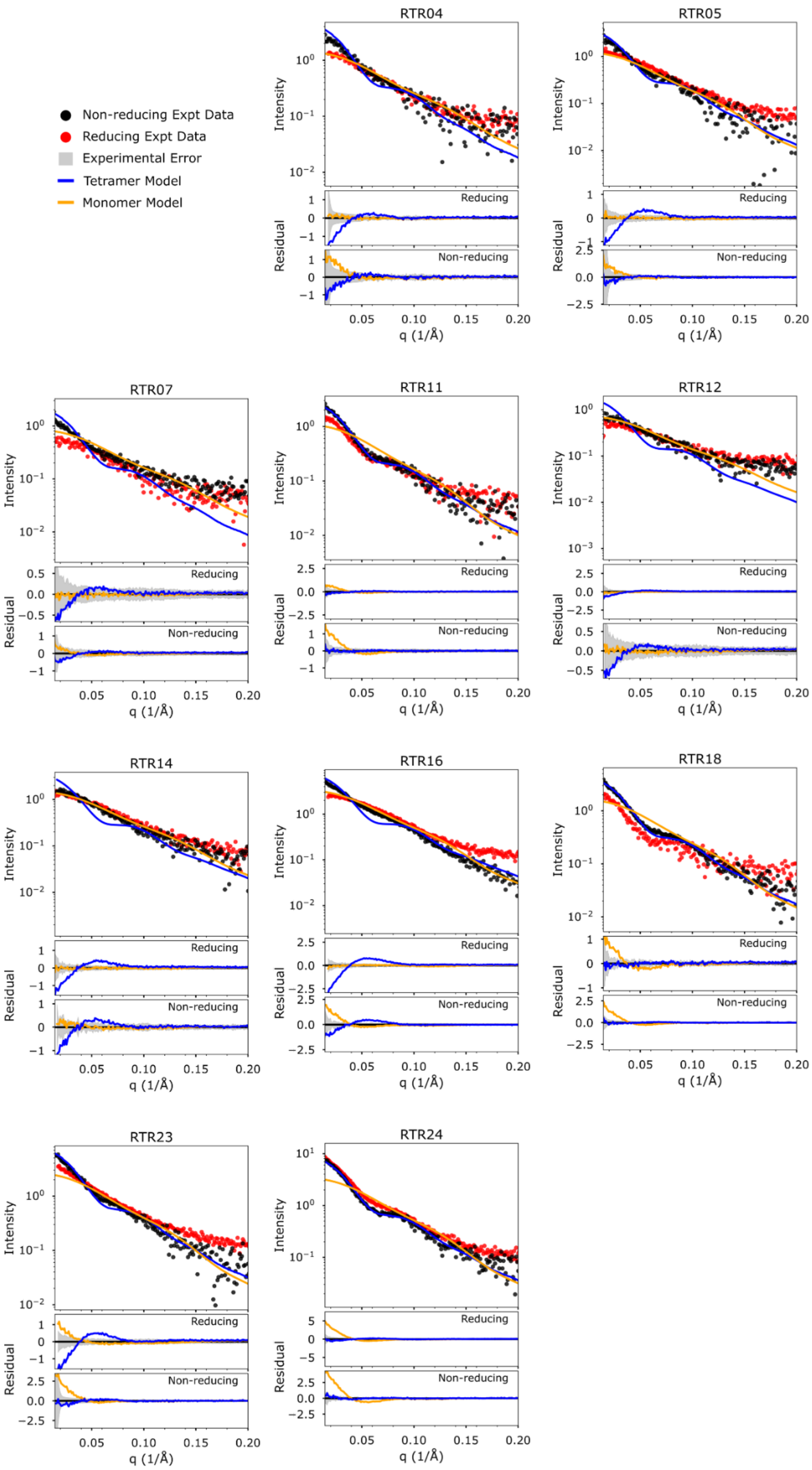

Non-reducing Expt Data
Reducing Expt Data
Experimental Error
Tetramer Model
Monomer Model
RTR04
RTR05
RTR07
RTR11
RTR12
RTR14
RTR16
RTR18
RTR23
RTR24
Intensity
Residual
Reducing
Non-reducing
q (1/Å)

**Supplemental Figure 15: SAXS of RTR homo-tetramers**

Plots of all designs with SAXS data. (Top of each panel) The scattering profiles plotting log intensity vs scattering angle. Black dots, averaged experimental data from many individual frames of non-reduced samples. Red dots, same as black dots but of reduced samples. Gray shading, the standard deviation of the averaged frames. Blue line, computed profiles generated from tetrameric design models and scaled using FoXS to fit the experimental data for the non-reduced data. Orange line, same as blue but with monomeric design models (Bottom of each panel) The residual of the fit between experimental data and the computed profile for reducing or non-reducing data, shown on a linear scale.

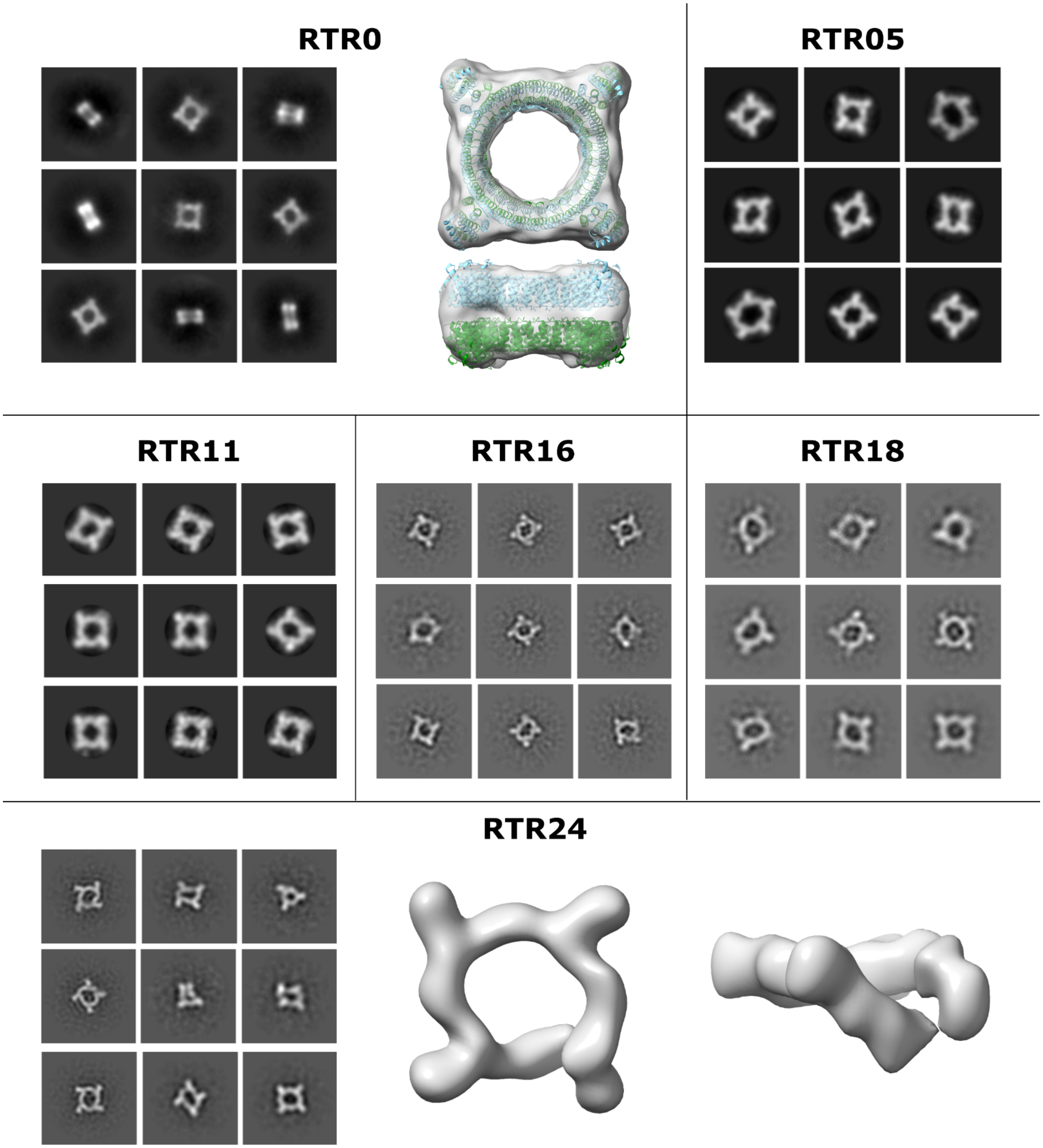


**Supplemental Figure 16: nsEM of RTR homo-tetramers**

Negative stain 2D class averages and 3D reconstructions of RTR0 and RTR variants, if possible. RTR16, RTR18, and RTR24 were processed without CTF correction during 2D classification. (RTR0) The map of RTR0 is an *ab inito* reconstruction with symmetry. Two copies of the design model of RTR0 were roughly fit into the map. (RTR24) The map of RTR24 is a heterogenous refinement without symmetry. Multiple classes of this map were observed with varying degrees of ring breakage.

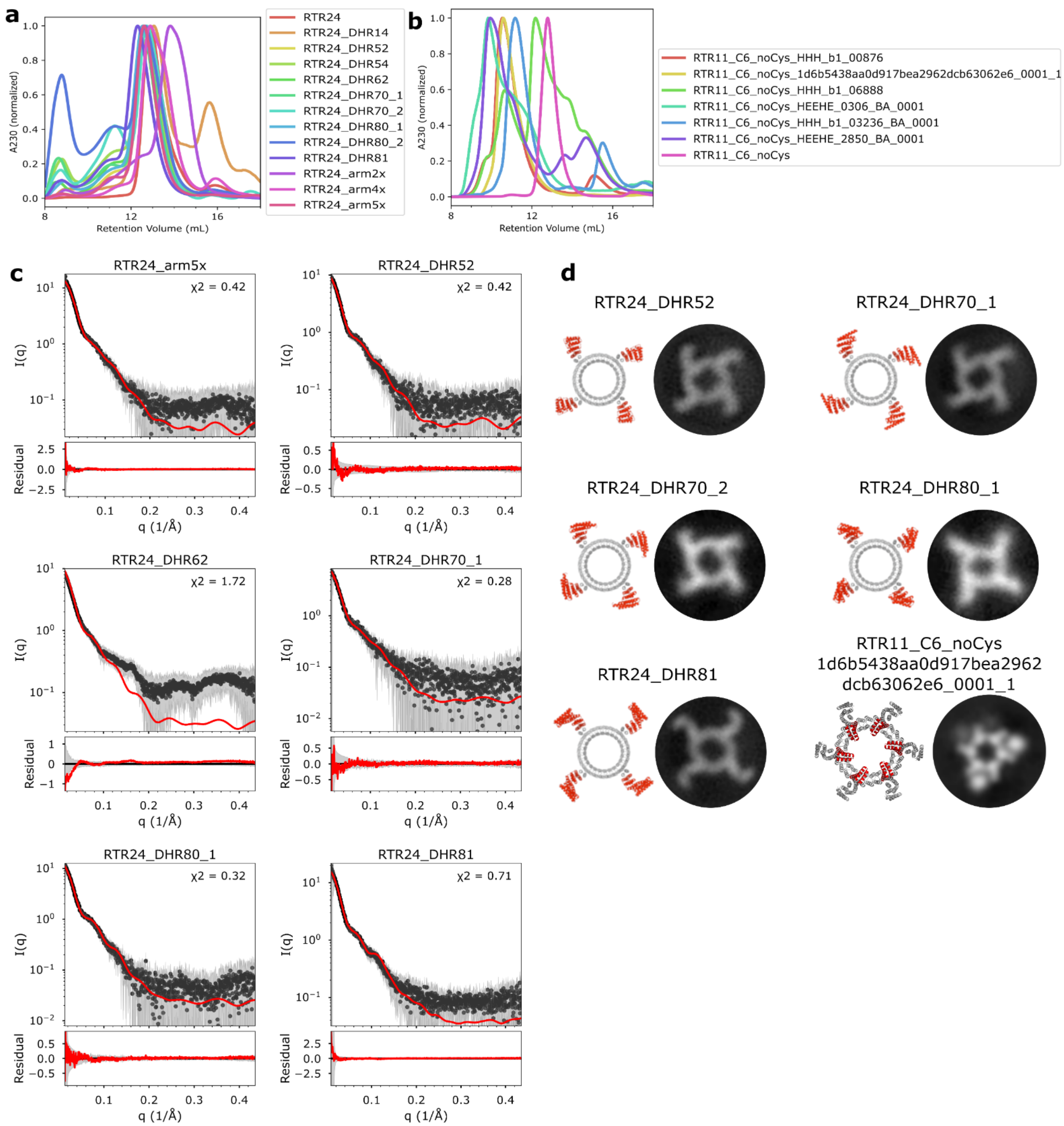

a
b
c
d
Retention Volume (mL)
A230 (normalized)
RTR24
RTR24_DHR14
RTR24_DHR52
RTR24_DHR54
RTR24_DHR62
RTR24_DHR70_1
RTR24_DHR70_2
RTR24_DHR80_1
RTR24_DHR80_2
RTR24_DHR81
RTR24_arm2x
RTR24_arm4x
RTR24_arm5x
RTR11_C6_noCys_HHH_b1_00876
RTR11_C6_noCys_1d6b5438aa0d917bea2962dcb63062e6_0001_1
RTR11_C6_noCys_HHH_b1_06888
RTR11_C6_noCys_HEEHE_0306_BA_0001
RTR11_C6_noCys_HHH_b1_03236_BA_0001
RTR11_C6_noCys_HEEHE_2850_BA_0001
RTR11_C6_noCys
RTR24_arm5x χ2 = 0.42
RTR24_DHR52 χ2 = 0.42
RTR24_DHR62 χ2 = 1.72
RTR24_DHR70_1 χ2 = 0.28
RTR24_DHR80_1 χ2 = 0.32
RTR24_DHR81 χ2 = 0.71
I(q)
Residual
q (1/Å)
RTR24_DHR52
RTR24_DHR70_1
RTR24_DHR70_2
RTR24_DHR80_1
RTR24_DHR81
RTR11_C6_noCys
1d6b5438aa0d917bea2962
dcb63062e6_0001_1

**Supplemental Figure 17: Characterization of RTR extensions**

(a) SEC traces of RTR24 with c-terminal extensions. RTR24 included as a size reference. (b) SEC traces of RTR11_C6 with n-terminal extensions. RTR11_C6 included as a size reference. (c) SAXS profiles of c-terminally extended designs. Black, experimental data. Red, theoretical profile based on design model fit to experimental data. Gray shading, standard deviation of averaged micrographs. (d) Negative stain EM 2D class averages of five c-terminally and one n-terminally extended RTR variants paired with cartoon models with the extended region shown in red. RTR11_C6_noCys_1d6b5438aa0d917bea2962dcb63062e6_0001_1 was tagged with GFP, which appear as ordered density between the outer arms.

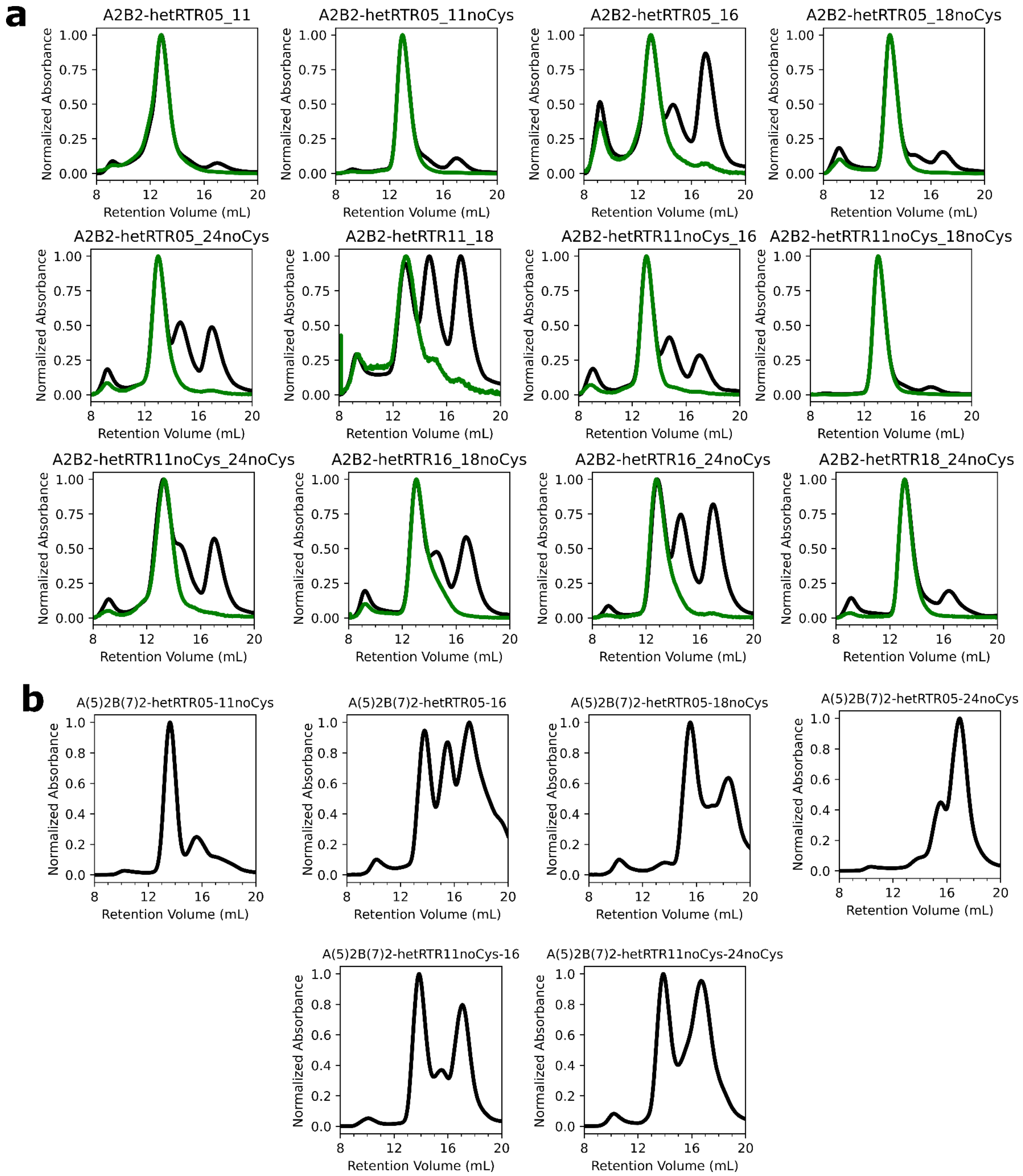


**Supplemental Figure 18: SEC of A2B2-hetRTRs**

(a) Black, normalized absorbance at 230nm. Green, normalized absorbance at 480nm, corresponding to the GFP-tagged chain. (b) Normalized absorbance at 230nm.

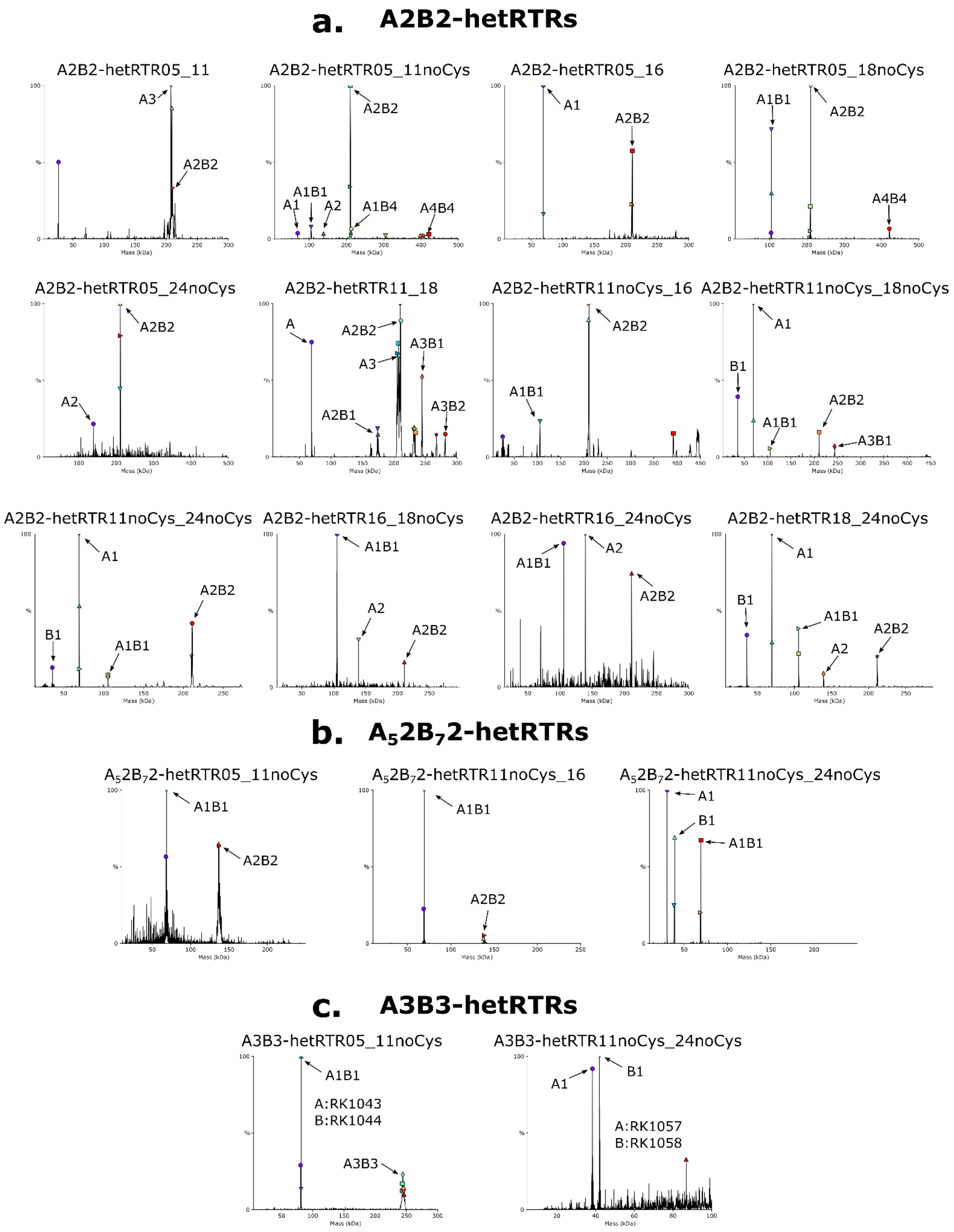


**Supplemental Figure 19: nMS of hetRTRs**

Plots of all designs of (a) A2B2-hetRTRs, (b) $A_52B_72$-hetRTRs, and (c) A3B3-hetRTRs with nMS data showing deconvoluted masses obtained from m/z spectra (see Supplemental Table 7). Identified species corresponding are annotated with arrows and labels. Unannotated peaks are unknown species.

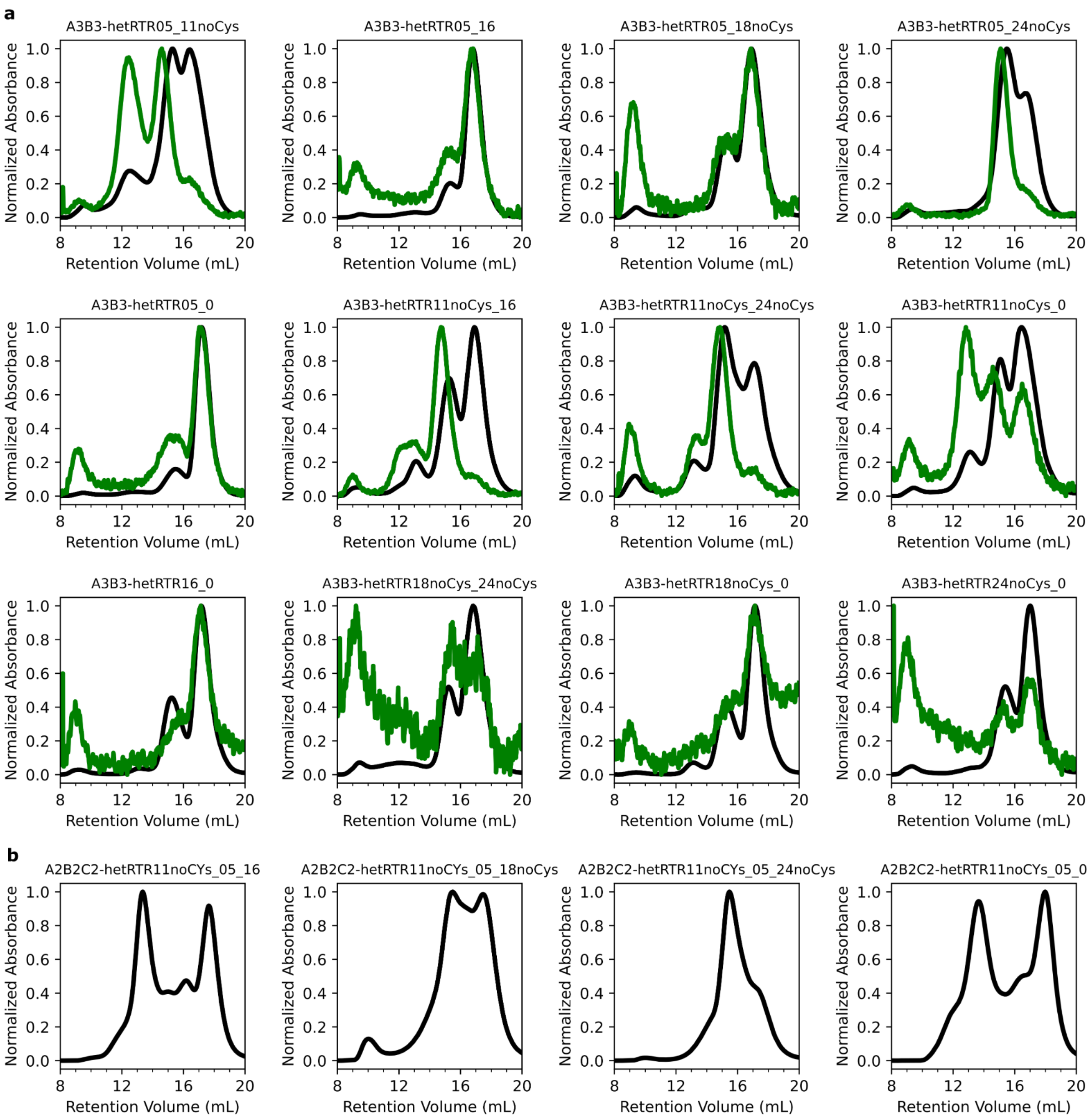


**Supplemental Figure 20: SEC of A3B3/A2B2C2-hetRTRs**

(a) Black, normalized absorbance at 230nm. Green, normalized absorbance at 480nm, corresponding to the GFP-tagged chain. (b) Normalized absorbance at 230nm.

**Supplemental Tables**

Supplemental Table 1: BGL homotrimer sequences
Supplemental Table 2: BGL crystallographic information
Supplemental Table 3: BGL heterotrimer sequences
Supplemental Table 4: RTR homo-oligomer sequences
Supplemental Table 5: A2B2-hetRTR sequences
Supplemental Table 6: A3B3/A2B2C2-hetRTR sequences
Supplemental Table 7: Native MS spectra of all designs

### Computational Methods

All code is available at https://github.com/rdkibler/pseudosymmetric_hetero-oligomers

While Rosetta is capable of performing every step of the interface redesign process using its symmetry machinery[1] and thus using a symmetric pose at every step, because we require the homology region to have identical sequence and coordinates between every design, we cannot make use of the concerted subunit rigid body transformation degrees of freedom that the symmetry machinery is, in part, intended to provide. Any full-chain movements away from the input structure in the homology region are not allowed. Additionally, we would not use the symmetry machinery's functions which make the packer and minimizer aware of the other chains in the structure due to the physical separation between the interfaces. Therefore, all steps until the final structure checks are performed on the structure asymmetrically using coordinate constraints to hold the homology regions in place. In the case of RTR0, we also got a large speed up by using a significantly truncated version of the interface region which included just the edge of the homology region.

#### 1. BGL redesign

All BGL design was carried out on the full 18 helix structure of BGL0 (PDB 6XR1), since this is semantically equivalent to working on the fixed-size homotrimer using symmetry operations, and avoids the need for symmetry which increases the computational overhead.

#### 1.1 Backbone perturbation

##### 1.1.1 Helix Rebuilding (HR) set

Canonically coiled alpha-helical backbones can be described by their Crick helical parameters[2,3], that is, for a given helix coiling around the Z-axis, the values of r0, omega0, omega1, phi0, phi1, detla_z, and whether the helix runs toward negative Z (inverted) or toward positive Z (not inverted). With the axis of structural symmetry lying on the Z-axis, the outer helices of BGL0 resemble the helix of a coiled coil with very large r0 (distance from the Z-axis). We sought to resample the backbone positions of the outer helix at the interface, and reasoned that replacement helices closeby in the parameter space of the original helix would also fit well physically while giving a large increase in backbone position (especially via sampling uniformly over phi1). To accomplish this, we wrote an algorithm which discovered the 7 helical parameters via least squares minimization. Most parameters were largely unconstrained save for r0 bounded between 1e-6 and 40 and phi0 bounded to >= 0. The parameters r0, omega0, omega1, phi0, phi1, and delta_z, were initialized to a generic left-handed supercoiled helix, with phi0 and r0 estimated from the position of the target helix's center of mass, and at every step the per-backbone-atom RMSD was computed between the target helix and a helix with the current parameters and a least squares algorithm was used to find the set of parameters which minimizes RMSD. The target in this campaign for BGL0 was the C-terminal helix, one of the outer ring of helices and the closest fit we achieved was 0.8A. Given this best description of the target helix as an ideal coiled coil helix, we deleted the target helix from the BGL0 structure and sampled 124,852 new helices with parameters r0 between 20 and 25, delta_z +/- 6 from the fitted value, omega0 +/- 1 degree from the fitted value, phi0 +/- 20 degrees from the fitted value, and phi1 +/- 154.3 degrees from the fitted value. Some solutions clashed with the rest of the BGL0 structure, so rather than fine-tuning parameter ranges which might have a complex relationship, we roughly evaluated clashing with a Rosetta scorefunction scoring only fa_rep, fa_atr, and fa_sol (delta REU from original structure < 9000) and buried surface area (at least 350 with either side of the interface).

The output of the helix rebuilding procedure is a disconnected helix roughly in the same position as the target helix and it needs to be re-connected to the rest of the structure via a loop. Because BGL0 is a toroid, the N- and C-termini are physically adjacent to each other and could be re-connected to either side. Therefore, for each output of helix rebuilding, we attempted to re-connect the helix as either the N-terminus (building a loop between the current N-terminus of the full structure and the new helix) or the C-terminus (building a loop between the current C-terminus of the full structure and the new helix) using ConnectChainsMover[4] with loop lengths between 2 and 5 residues and the option to add or remove five residues from either side of the loop. This resulted in the creation of 3524 outputs each for N-term closure and C-term closure.

*Extra functionality*

While not necessary for this work, we found that Crick helical parameters do not always generalize to helical segments which appear curved like coiled coils but, for example, whose superhelical curvature is on an axis which does not pass through the center of mass of the protein. To account for this, we have also developed a method for fitting a reduced set of Crick helical parameters and 6 additional parameters corresponding to a homogeneous transform (3 rotation and 3 translation) of the helix using least squares minimization. The parameters made redundant by this approach and which are thus never sampled are delta_z and invert because they can be easily described by the transform parameters. The algorithm proceeds in three phases.

In the first phase, the goal is to roughly locate the superhelical axis. The helix parameters are initialized to those of a generic left-handed superhelical helix (r0 = 5, phi1=0 degrees, phi0=100 degrees, omega0=-2.85 degrees, omega1=102.85 degrees) and the transform is initialized to a rough initial guess based on the target helix's center of mass and orientation of the termini relative to the Z-axis. Phi1 and transform are fitted to the target helix to refine the initial guess of the transform of the superhelical axis. The signal from the phase of the minor helix (the atomic positions) is much stronger than the major helix (superhelical axis), so phi1 is allowed to vary to avoid biasing the transform by a bad guess of phi1.

In the second phase, which may not be necessary, the transform is fixed and omega1, omega0 (coupled to omega1 via the relationship abs(abs(omega0+omega1)-radians(100)) < 10), phi0, and phi1 are allowed to minimize in order to roughly shape the helix and get it close to its final form in the context of a new superhelical axis.

In the third phase, r0, omega0, omega1, phi1 (not phi0), and the transform are allowed to vary to simultaneously fine-tune the helical parameters and the effective location of the superhelical axis. The tolerance of the fit is set much tighter than for phases 1 or 2. Given a set of parameters that describe the new helix, a similar closeby parameter search and hash table clash check could be performed.

**1.1.2 Normal Modes (NM) Relaxation**

Separately from HR, we generated conformations very similar to the native structure but which may differ only via molecular breathing type motions by applying the NormalModeRelaxMover (NMRM)[5] to generate another set of designs (NM). We restricted backbone perturbation to the two and a half helices on either side of the interface (N-term Interface: n-term through the middle of the 3rd helix; C-term Interface: the middle of the 3rd-to-last helix through the c-term), and additionally used coordinate constraints to hold everything else in place. NMRM was run successively and separately on the C-term Interface then the N-term interface, with the rationale that allowing the movement of the C-term Interface and its outer C-terminal helix first would create room for a greater range of movement by the N-term Interface. 500 backbones were generated with pertscale=2.0, nmodes=60, and mix_modes=true for both NMRM runs.

**1.2 HBNet design**

HBNet design was carried out slightly differently depending on the design set. Common to both were the region within which HBNets could be designed (the union of the N-term Interface and C-term interface), as well as a requirement to have at least 4 network residues in the core, have no unsatisfied heavy atoms or polar hydrogens, make at least 2 intermolecular hbonds, and have between 1 and 3 networks in the output structure. Hb_threshold was set to -0.6 and the MC sampling was turned on with seed_hbond_threshold -0.4 and 10,000 MC steps for HR and 100,000 for NM for deeper sampling. Core cutoff for HR was set to SASA < 20 (ball radius 2.5), but for HR was SASA < 6 (more stringent) to encourage deeper HBNets at a cost to sampling.

The HBNets created for the NM set were highly redundant due to the similarity of the backbones inputs and depth of sampling, so the 12954 outputs were grouped by the sequence of their buried HBNet residues (SASA < 20 with ball radius 2.5) and the best scoring output was kept for each group (3332 total).

The HBNets for HR could not be grouped as easily but were qualitatively more diverse, so the 112039 outputs were instead briefly fixed-backbone designed with HBNet's AtomPair distance constraints on and sidechain relaxed with constraints off to allow poorly positioned and/or poorly packed HBNets to move out of position. Then, designs were filtered on vbuns <= 2, # alanines <= 126, and total HBNet constraint score (indicating movement of HBNet residues) < 1.0, yielding 2538 outputs. The new helix and new loops were cartesian minimized in preparation for interface design.

#### 1.3 Interface design

In both cases, flexible backbone FastDesign was performed with the standard suite of TaskOperations (InitializeFromCommandline, DesignRestrictions, IncludeCurrent, LimitAromaChi2, and ExtraRotamersGeneric [ex1 & ex2]). Design was carried out with variable bondangle and bondlengths, and the approximate_buried_unsat_pentality and PruneBuriedUnsats[6] were used to aid in retaining or extending the HBNets. Flexible backbone movement was only allowed on the N-term Interface and C-term Interface regions, but sequence design was allowed at the interface between the Interface regions and in a 7A neighborhood around each HBNet residue (but not including the HBNet residues or proline or glycine). The residues within 6A of the designable region were allowed to repack and sidechain minimize and the HBNet residues had their AtomPair constraints enabled. FastDesign was carried out with a modified version of the MonomerDesign2019 relax script which does not ramp coordinate constraints.

##### 1.3.1 HR specific

Calpha atoms were loosely coordinate constrained over the entire pose. The pose has a reasonable sequence already from the HBNet filtering step, so only minor changes to core packing were allowed for L, I, and V residues which were only allowed to change to L, I, or V. PruneBadRotamers and ConsensusLoopDesign TaskOperations were also employed. The surface was not designed in order to speed up design around the HBNets which were buried. Finally, because the new helix created by HR was given a “rosetta default” sequence, additional amino acid composition constraints were used to constrain the structure to have an amino acid sequence profile roughly similar to the original design, namely requiring at most 1 methionine, 127 alanines, 2 phenylalanines, and 5 threonines, with some combination of linearly increasing penalties within a small range then increasing quadratically at higher counts. 7 separate design trajectories were run on each input, resulting in creation of 12256 outputs.

**1.3.2 NM specific**

Calpha atoms were tightly coordinate constrained only over the non-Interface region to allow more movement at the Interface. 7 separate design trajectories were run on each input, resulting in creation of 12879 outputs.

**1.4 Filtering**

Designs were filtered on a variety of criteria to reduce each design set to 100 for manual inspection. The criteria are the following:

HR:

- Cst_after < 1.0
- Mismatch_probability < 0.06
- Network_holes < -1.4
- Interface_holes < 0
- dSASA_polar > 1500
- vbuns5.5_heavy_ball_1.1D = 0
- RotamerBoltzmannWeight (HBNet resis) < -0.51
- lowest 100 by worst9mer

NM:

- Cst_after <1.0
- Network_holes < -1.4
- Interface_holes < 0
- P_aa_pp < -130
- Mismatch_probability < 0.07
- vbuns5.5_heavy_ball_1.1D = 0
- RotamerBoltzmannWeight (HBNet resis) < -0.52
- Highest 100 by dSASA_polar

The sets were then manually inspected, visually looking for hydrophobic interdigitation in the core, interesting and practical HBNets, and reasonable burial of HBNets. Finally, 16 HR designs and 4 NM designs from each set of 100 were rearranged to be (6 helix) homotrimers, ordered, and characterized.

**1.2 RTR redesign**

Design was carried out on a truncated sub-structure of RTR0 focusing on the interface region, comprising just 210 residues which represent four helices from the N-terminal side of the interface (two participating in the interface and two representing the fixed homology region) and 8 helices from the C-terminal side of the interface (4 which participate in the interface, two which reprepresent the fixed homology region of the ring, and two which represent the fixed part of the C-terminal extension).

1.1 Backbone perturbation

NMRM parameters were optimized to create a variety of pocket sizes and shapes in the interface (pertscale = 1.6, nmodes = 31, mix_modes=true). The regions allowed to move, Interface A and Interface B, were defined as regions 1-33 and 100-174 on the truncated “mini”-pose RC4_20_mini.pdb. Before running NMRM, the sequence of the pose was changed to polyvaline and the beta_nov16_soft weights were used in order to allow the structures to slide past each other more easily. NMRM was also run in a single step, letting both interfaces move at once. After perturbation, the original sequence was restored. A total of 15500 outputs were produced.

To reduce the runtime of the protocol, the NMRM source code (source/src/protocols/normalmode/NormalModeRelaxMover.cc) was modified. NMRM stores all computed conformations it creates during the course of a given trajectory but only returns the best scoring one (throwing away the rest). We updated the mover to allow retrieval of all computed conformations via a MultiplePoseMover in either a random or best-score-sorted order.

**1.2 HBNet design**

The region where HBNets can be designed into the RTR interface is smaller and empirically more difficult to find extensive HBNets at diverse positions. Therefore, HBNet design in the case of RTR was modified to require fewer HBNet residues to be in the core (2) and filtering was set up to require that at least one buried HBNet residue be on either side of the interface. max_unsat_Hpol was also increased to two. See production_varset.json for a complete list of parameters used.

HBNet produced 15692 outputs and we found 3815 unique networks. We kept either the best 3 or the best 10% of structures with each network, whichever was higher, for a total of 12649.

**1.3 Interface design**

The interface between Chain A and Chain B and the region 7.0 A around the HBNet residues, not including the HBNet residues themselves, were allowed to redesign, excluding serine 154 which seemed to be important. Flexible backbone movement was allowed for the n-terminal two helices and loops on chain A and the four helices which are closest to chain A on chain B with all other residues (essentially the two helices on chain A and chain B which connect the interface to the rest of the ring and the two helices which connect the interface to the rest of the extension on chain B) were fixed in place with coordinate constraints and prevented from backbone minimization. Then flexible backbone FastDesign was carried out in cartesian space with HBNet residue AtomPair distance constraints, allowing at most 2 methionines, allowing at most 2 boundary tryptophans, and requiring between 1 and 3 boundary tyrosines. The same task operations were used as 1.3 for NM designs.

Next, the surface was separately redesigned with sequence constraints which require a very negative net charge to ensure any combination of interfaces downstream has a very low isoelectric point (pI). Designs were filtered to have mismatch_probability, core_holes, interface_sc, and ss_sc not worse than the parent structure, resulting in 880 unique outputs.

The minimal structures were then converted back into the full (tetrameric) ring through superimposition of the homology region helices with the corresponding regions of the full structure and splicing of the redesigned interface into the full structure using PyMol (Schrödinger). The symmetric structures were then cartesian minimized with HBNet constraints on with Rosetta to fix any bad backbone geometry from the splicing procedure, and finally they were flexible backbone relaxed without HBNet constraints.

#### 1.4 Filtering

DeepAccNet[7] was used to predict the lddt of the designs post-relaxation, and any designs with residues with lddt < 90 were discarded. Designs which did not increase much in energy after flexible backbone relaxation without HBNet constraints (REU after - REU before < 0.5) were also discarded, resulting in 79 designs which were subsequently visually inspected and narrowed down to 24 designs.

All interface design was previously carried out while disallowing cystine, but an interface disulfide seemed to have been important for high order structure assembly[8], so we used stapler (https://github.com/atom-moyer/stapler) to consider potential positions for disulfide bonds to exist on the selected designs, then chose positions which were as different as possible from the others in the chosen set. These were then ordered and characterized.

### 2. Computational recombination

#### 2.1 hetBGLs

The process of combining multiple homo-oligomers into one hetero-oligomeric structure was carried out using a python script, recombine_2.1.py. It uses PyMol to superimpose and splice together homo-oligomer structures at the specified recombination point in the desired order, controlled through commandline arguments. The recombined structure could then be directly outputted, or optimized using Rosetta cartesian backbone minimization to fix any bad backbone geometry. Additionally, because local interface pI was not considered during the initial BGL design, some naive combinations could have individual chains with pI >= 8, which would result in neutrally or positively charged chains which are more likely to have expression and stability issues. Therefore, the surfaces of hybrid chains could also be redesigned to enforce a small pI for each chain individually in the final hetero-oligomers. Finally, if not present before, tyrosines were added to the surfaces to act as crystal contacts to aid crystallography.

**2.2 hetRTRs**

Recombination of RTRs into hetRTRs was done either using a version of the recombinase script which does not perform surface design, or with a text editor to simply combine the amino acid sequences to form the appropriate hybrid chains. In the latter case, design models were constructed manually using PyMol.

**Experimental Methods**

**Buffer and media recipes**

All buffers and media were made using Milli-Q filtered water.

*Autoinduction media (TBM-5052)*

1.2% [wt/vol] tryptone, 2.4% [wt/vol] yeast extract, 0.5% [wt/vol] glycerol, 0.05% [wt/vol] D-glucose, 0.2% [wt/vol] D-lactose, 25 mM Na2HPO4, 25 mM KH2PO4, 50 mM NH4Cl, 5 mM Na2SO4, 2 mM MgSO4, 10 µM FeCl3, 4 µM CaCl2, 2 µM MnCl2, 2 µM ZnSO4, 400 nM CoCl2, 400 nM NiCl2, 400 nM CuCl2, 400 nM Na2MoO4, 400 nM Na2SeO3, 400 nM H3BO3

*Lysis buffer*

25 mM Tris, 300 mM NaCl, 20 mM imidazole, 10% glycerol, pH 8.0 at room temperature

*Wash buffer*

25 mM Tris, 300 mM NaCl, 40 mM imidazole, 10% glycerol, pH 8.0 at room temperature

*Elution buffer*

25 mM Tris, 300 mM NaCl, 300 mM imidazole, 100 mM EDTA, 10% glycerol, pH 8.0 at room temperature

*Het Lysis buffer*

25 mM Tris, 300 mM NaCl, 10 mM imidazole, 10% glycerol, pH 8.0 at room temperature

*Het Wash buffer*

25 mM Tris, 300 mM NaCl, 20 mM imidazole, 10% glycerol, pH 8.0 at room temperature

*Het Elution buffer*

25 mM Tris, 300 mM NaCl, 50 mM imidazole, 10% glycerol, pH 8.0 at room temperature

*SEC buffer*
25 mM Tris, 300 mM NaCl, pH 8.0 at room temperature

*SAXS buffer*
25 mM Tris, 300 mM NaCl, 2% glycerols, pH 8.0 at room temperature

*TEV buffer*
25 mM Tris, 100 mM NaCl, 0.5 mM EDTA, 1mM DTT, pH 8.0 at room temperature

*AEC buffer A*
25 mM Tris, pH 8 at room temperature

*AEC buffer B*
25 mM Tris, 1M NaCl, pH 8 at room temperature

**Construction of synthetic genes**

All synthetic genes were ordered from either Integrated DNA Technologies Inc. (Coralville, IA, USA) (IDT) or Genscript Inc. (Piscataway, NJ, USA). Genes ordered from IDT were reverse translated and codon optimized using Domesticator (https://github.com/rdkibler/domesticator_3). Protein tags were added to aid in purification (His6-tag) or identification (EHEE_rd2_0005 (aka EHEE)[9], superfolderGFP, or mScarlet-I) were separated from the designed protein with a tobacco etch virus protease (TEVp) cleavage site (ENLYFQG). In many cases, stop codons were introduced at the end of the inserted gene to prevent incorporation of the vector's built-in tags. Cloning into pET29b+ or its derivatives is always done at the NdeI/NcoI site.

For expression of BGLs, protomer sequences were extracted and the n-terminal tag "MGHHHHHHGSENLYFQGWS" was added. Codon-optimized genes with a 3' stop codon were cloned into pET29b+ by IDT. See Supplemental Table 1.1 for full amino acid sequences.

For co-expression of hetBGLs, all three proteins were arranged sequentially off the same transcript, separated by ribosome binding sequences (RBSs) (first: TAAGAAGGAGATATCATCATG; second: TAAAGAAGGAGATATCATATG). One protomer received superfolderGFP, another received EHEE, and the third received nothing. One protomer (with either superfolderGFP for the EHEE) received a His6-tag. See Supplemental Table 1.3 for full amino acid sequences. These were ordered from IDT.

For expression of RTRs, amino acid sequences from Supplemental Table 1.4 were reverse translated and inserted into pET29b+ with a stop codon added before the end of the insert. These were ordered from IDT.

For co-expression of tetrameric hetRTRs, amino acid sequences from Supplemental Table 1.5 corresponding with the A2B2-hetRTRs were reverse translated and inserted into a pET29b+ vector variant which expresses the n-terminal superfolderGFP tag such that chain A was tagged with TEV-cleavable sfGFP. Chain A and B were separated by an RBS (TAAGAAGGAGATATCATCATG), and chain B received the vector's c-terminal His6-tag. The sequences corresponding to A(5)2B(7)2 were prepared in the same way but inserted into the standard pET29b+ vector. All C4 sequences were ordered from IDT and everything else was ordered from Genscript.

For (separate) expression of hexameric hetRTRs, the amino acid sequences from Supplemental Table 1.6 were reverse translated and inserted into pET29b+ using the vector's c-terminal His6-tag. These were ordered from Genscript.

**Protein production**

**Transformation and expression**

For single plasmid expressions, plasmids (100ng) were transformed into chemically competent E. coli expression strain BL21(DE3)Star (Invitrogen) for protein expression following manufacturer's protocol, with the exception of using 10ul competent cells per reaction. Following transformation and recovery, the entire transformation products were used to inoculate 1 mL Luria-Bertani (LB) medium containing 100 ug/mL kanamycin and grown at 37°C with shaking at 225 rpm overnight. 500ul of overnight cultures were diluted into 50 mL TBM-5052 supplemented with 100 ug/mL kanamycin in 250 mL baffled flasks, and incubated at 37°C with shaking at 225 rpm for 18-24 hours.

**Immobilized metal affinity chromatography (IMAC)**

Cultures were harvested by centrifugation at 4000 rcf for 10 minutes, culture supernatant decanted, and pellets resuspended to 30 mL in Lysis buffer. 300ul PMSF (100mM in 100% EtOH) is added immediately prior to sonication at 70% power for 5 minutes. "Lysate" fractions are saved, and then lysates were clarified by ultracentrifugation at 14,000 rcf at 12°C for at least 30 minutes and applied to 1.5 mL Ni-NTA resin (Qiagen) pre-equilibrated with Lysis buffer and packed into Econo-Pac columns (Bio-Rad) for gravity chromatography. The columns were washed twice with 15 mL Wash buffer and eluted with 10 mL Elution buffer.

Co-expressed hetero-oligomers were purified according to a similar procedure, except they used the “Het” variants of the the Lysis, Wash, and Elution buffers, followed by a final elution with the standard Elution buffer, as this was found to improve the yield of single-His6-tagged complexes over non-specific multiple-His6-tagged complexes which can arise at high concentrations and likely dominate binding to the Ni-NTA resin. Samples prepared for crystallization were treated similarly, except 500 mL cultures were used and lysate was divided among six gravity columns.

**Size-exclusion chromatography (SEC)**

Samples were concentrated using 10k MWCO spin concentrators and were purified using a Superdex 200 10/300 increase column (Cytiva) in SEC buffer using an ÄKTA pure system (Cytiva). SEC traces were also used to qualitatively determine homogeneity and quantitatively measure total yield by A280 absorbance integrated over the collected fractions using Unicorn (Cytiva).

**TEVp cleavage**

Purification and mass tags were buffer exchanged into TEV buffer and cleaved with TEVp at a ratio of 1 mg TEV per 100 mg substrate for 24-72h at room temperature. After TEV cleavage, samples were exchanged into Lysis buffer and passed over a Ni-NTA gravity column and washed with 10ml lysis buffer. Flowthrough was collected, concentrated using 10k MWCO spin concentrators, and purified once again by SEC.

**Anion exchange chromatography (AEC)**

TEVp-cleaved samples of hetBGL03-15-18 intended for crystallization remained contaminated with GFP due to an oversight that meant GFP did not have a His6-tag. To remove the GFP, samples were exchanged into AEC buffer A and loaded onto a HiScreen Q FF (Cytiva) column pre-equilibrated in AEC buffer A using an ÄKTA pure system. A gradient of AEC buffer B was applied with pauses at 15% (150 mM NaCl) and 25% (250mM NaCl) to elute the GFP and hetBGL03-15-18, respectively. Separation was measured by differential absorbance at 480 nm (GFP absorbance) and 280 nm, and SDS-PAGE.

**Sample analysis**

**SDS-PAGE**

Samples were diluted 1:1 with 2x Laemmli Sample Buffer (Bio-Rad) without Beta-mercaptoethanol and 15ul were loaded onto AnykD™ Criterion™ TGX™ Precast Midi Protein Gels (Bio-Rad). Ladder was 10ul of Precision Plus Protein™ Kaleidoscope™ Prestained Protein Standards (Bio-Rad). Gels were run at 300V for 18 minutes, then stained using an eStain™ L1 Protein Staining System (Genscript). Stained gels were imaged using a Chemidoc XRS+ (Bio-Rad)

**Liquid chromatography mass spectrometry (LC-MS)**

To identify the molecular mass of each protein and thus verify sample identity and integrity, intact mass spectra was obtained via reverse-phase LC/MS on an Agilent G6230B TOF on an AdvanceBio RP-Desalting column, and subsequently deconvoluted by way of Bioconfirm using a total entropy algorithm.

**Native mass spectrometry (nMS)**

Online buffer exchange coupled to native mass spectrometry (OBE-nMS) was used to analyze the oligomeric states of SEC-purified and LC-MS verified samples. Using a Vanquish Duo Ultra-High-Performance LC (UHPLC) system (Thermo Scientific) coupled to a Q Exactive Ultra High Mass Range (UHMR) Hybrid Quadrupole-Orbitrap mass spectrometer (Thermo Scientific) and desalting cartridge prototypes (~2 mm x 25, 30, or 50 mm; 80, 120, or 250 Å pore size; packed with 3 µm silica or polymer particles) provided by Thermo Fisher Scientific (Sunnyvale, CA), samples were buffer exchanged into 200 mM ammonium acetate[10,11]. Mass spectra were then deconvolved with UniDecs versions 4.1.2 and later[12].

**Size Exclusion Chromatography - Multi Angle Light Scattering (SEC-MALS)**

IMAC and SEC purified samples were were analyzed by SEC-MALS in 20 mM Tris, 150 mM NaCl, pH 8 on a Superdex 200 10/300 column in line with a Heleos multi-angle 9 static light scattering and an Optilab T-rEX detector (Wyatt Technology Corporation). Data was analyzed using ASTRA (Wyatt Technologies) to calculate the weighted average molar mass (Mw) of the selected species.

**Negative stain electron microscopy (nsEM)**

SEC purified samples were diluted to ~0.005 mg/ml using SEC buffer immediately before application for 45s to glow discharged thick carbon film-coated 400 mesh copper grids (CF400-CU TH) (Electron Microscopy Sciences). Grids were then stained and dried immediately twice using 2% uranyl formate. Dried grids were screened on a 120 kV Talos L120C transmission electron microscope. The *E. Pluribus Unum* (EPU) (FEI Thermo Scientific) software was used for automated data collection. Data processing was carried out in CryoSPARC™ (Structura Biotechnology Inc).

**Small angle X-ray scattering (SAXS)**

TEVp-cleaved (and optionally AEC purified) samples were re-purified by SEC in SAXS buffer and concentrated using thoroughly washed 10k MWCO small spin concentrators; the flowthrough of concentration was used as blanks for buffer subtraction. Scattering measurements were performed at the SIBYLS 12.3.1 beamline at the Advanced Light Source as part of the HT-SAXS program. The Xray wavelength ($\lambda$) was 1.27 Å, and the sample-to-detector distance was 1.5 m, corresponding to a scattering vector q ($q = 4\pi \sin \theta/\lambda$, where $2\theta$ is the scattering angle) range of 0.01 to 0.3 $Å^{-1}$. A series of exposures, in equal subsecond time slices, were taken of each well: 0.3 second exposures for 10 seconds resulting in 32 frames per sample. For each sample, data was collected for two different concentrations to test for concentration dependent effects; "low" concentration samples were ~2.5 mg/mL and "high" concentration samples were ~ 5 mg/mL. Data was processed using the SAXS FrameSlice online server (https://bl1231.als.lbl.gov/ran). FoXS[13] was used to compare design models to experimental scattering profiles and calculate quality of fit ($\chi$) values. The SAXS Similarity online server was used to compute the similarities of scattering profiles to each other and calculate quality of fit ($\chi$) values.

**X-ray crystallographic analyses**

Crystals of BGL06, BGL14_styr, BGL15, BGL18, and hetBGL03-15-18 were grown using protein purified as described above and TEVp cleaved and optionally AEC purified. Protein samples dispensed in 1 uL drops at purification concentrations were mixed with equal volume of a crystallization solution and set in hanging drops (refer to Supplemental Table 1.2 for conditions). Vapor phase equilibration of the resulting drops against a 1 mL reservoir of the same crystallization solution resulting in growth of crystals. The crystals were flash cooled in liquid nitrogen after transfer into a cryoprotective solution (refer to Supplemental Table 1.2 for conditions). Diffraction data were collected on a Pilatus areas detector at the Advanced Light Source (ALS) synchrotron facility at beamline 5.0.2 for BGL06, BGL14_styr, BGL15, and BGL18. Diffraction data were collected on a Rigaku HyPix-6000HE hybrid photon counting detector at the Fred Hutchinson Cancer Center (FHCC) for hetBGL03-15-18. The resulting data sets (Supplemental Table 1.2) extend to 2.1 Å, 3.0 Å, 3.3 Å, 3.0 Å, and 2.1 Å resolution for BGL06, BGL14_styr, BGL15, BGL18, and hetBGL03-15-18, respectively. Most data had complete trimers within the asymmetric unit (three copies of a protein subunit), with exceptions to BGL18 and BGL15 which had 2 and 4 trimers in the asymmetric unit, respectively. Data was processed using the program HKL2000[14] or Aimless[15]. The placement of subunits was determined using the molecular replacement algorithm in program PHENIX[16]. Local rebuilding of all constructs was performed using the program COOT[17] , followed by refinement using the program PHENIX[16]. The final values for Rwork / Rfree are notated in Supplemental Table 1.2.
